# Biophysical model for predicting muscle short-range stiffness during movement

**DOI:** 10.1101/2025.10.31.685881

**Authors:** Tim J. van der Zee, Surabhi N. Simha, Gregory N. Milburn, Kenneth S. Campbell, Lena H. Ting, Friedl De Groote

## Abstract

Musculoskeletal simulations can offer valuable insight into how the properties of our musculoskeletal system influence the biomechanics of our daily movements. One such property is muscle’s history-dependent initial resistance to stretch, also known as short-range stiffness, which is key to stabilizing movements in response to external perturbations. Short-range stiffness is poorly captured by existing musculoskeletal simulations since they employ phenomenological Hill-type muscle models that lack the mechanisms underlying short-range stiffness. While it has been previously shown that biophysical cross-bridge models can reproduce muscle short-range-stiffness, it is unclear which specific biophysical properties are necessary to capture history-dependent muscle force responses in behaviorally relevant conditions. Here, we tested the ability of various biophysical cross-bridge models to reproduce empirical short-range stiffness and its history-dependent changes across a broad range of behaviorally relevant length changes and activation levels, using an existing dataset on permeabilized rat soleus muscle fibers (N = 11). We found that a biophysical cross-bridge model with cooperative myofilament activation reproduced the effects of muscle activation (R^2^ = 0.86), stretch amplitude (R^2^ = 0.71) and isometric recovery time (R^2^ = 0.79) on history-dependent changes in short-range stiffness after shortening. Similar results were obtained when the cross-bridge distribution of the biophysical model was approximated by a Gaussian (R^2^ = 0.73 - 0.88), but at a 20 times lower computational cost. These effects could not be reproduced by either a biophysical cross-bridge model without cooperative myofilament activation or a Hill-type model (R^2^ < 0.5). The reduced computational demand of the Gaussian-approximated models facilitates implementing biophysical cross-bridge models with cooperative myofilament activation in musculoskeletal simulations to improve the prediction of short-range stiffness during movements.

## Introduction

Skeletal muscles are both the motors and the brakes of our musculoskeletal system (Dickinson et al., 2000). Muscles’ ability to act as a brake is especially important for stabilizing movements in response to perturbations, reducing the need for sensorimotor feedback (Brown and Loeb, 2000; Daley and Biewener, 2006; Loeb, 1995; Ting et al., 2009). Short-range stiffness, a rapid and transient rise in muscle force upon stretch, is a movement history-dependent muscle property that contributes to a muscle’s braking ability (Rack and Westbury, 1974; Campbell and Moss, 2000). Most of this rapid increase in force is thought to arise from cross-bridge stretching, i.e. cross-bridges act as stiff springs (Fig. 1). Prior muscle shortening results in a subsequent reduction in short-range stiffness that is known as muscle thixotropy. This stiffness reduction after shortening is consistent with a cross-bridge mechanism, as shortening can reduce the number of attached cross-bridges (Fig. 1). Indeed, biophysical models of cross-bridge cycling can qualitatively reproduce short-range stiffness and muscle thixotropy (Campbell and Moss, 2002), while phenomenological Hill-type models cannot (De Groote et al., 2017; Willaert et al., 2024). However, simple cross-bridge models may be insufficient to capture both the stretch and shortening dynamics of muscle (Cole et al., 1996; Lakie and Campbell, 2019; Simha and Ting, 2024; van den Bogert et al., 1998). We therefore aimed to identify a biophysical muscle model that could capture behaviorally-relevant muscle length changes, including short-range stiffness and its history dependence at a range of activation levels measured in vitro (Horslen et al 2023). As cross-bridge models are more computationally intensive than Hill-type models, we also sought to improve the computational efficiency of the resulting model.

**Figure 1.**
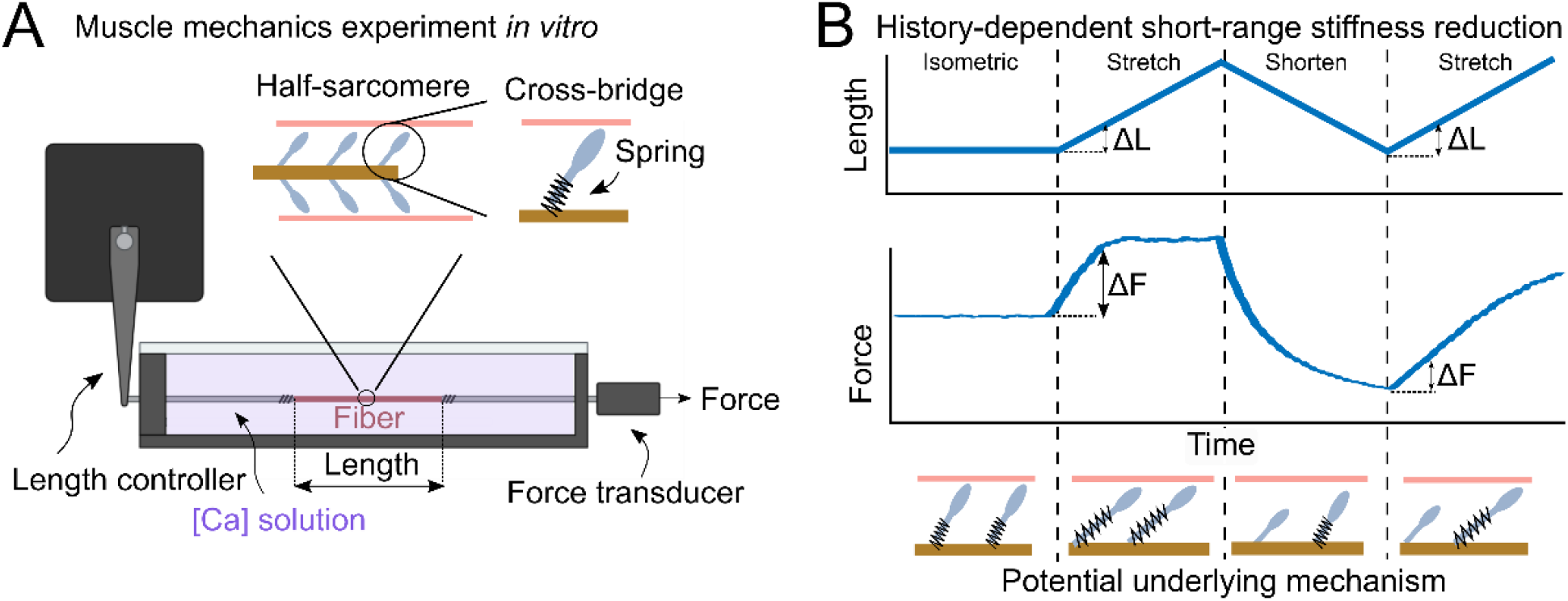
Muscle thixotropy is a history-dependent stiffness reduction. **A**. Experimental set-up commonly used for characterizing muscle short-range stiffness. A single permeabilized muscle fiber is attached to a length controller on one end and a force transducer at the other. Muscle activation is controlled using a bathing solution with a fixed calcium concentration. **B**. Typical example data from a muscle mechanics experiment in which triangular length changes are imposed on an isolated muscle fiber. Force rapidly increases during the first stretch (Δ*F*). But when the fiber is shortened and stretched again, the force increase Δ*F* is less for the same length increase Δ*L*. This may be explained by molecular cross-bridges, which act as springs when attached but detach during shortening. Cross-bridges may remain detached for some time after shortening, potentially explaining why force increases less rapidly during the second stretch compared to during the first stretch. Part of the figure was designed using BioRender.

**Movement history-dependent changes in muscle short-range stiffness known as muscle thixotropy may result from cooperative myofilament dynamics that regulate cross-bridge cycling**. Cross-bridge cycling is a biomolecular reaction, where the rate of cross-bridge attachment is the product of a rate constant, the number of available myosin heads, and the number of available thin filament binding sites. Cross-bridge attachment is cooperative in at least three ways. First, the activation of the thin filament (i.e., the fraction of actin binding sites that can bind cross-bridges) increases as the number of attached cross-bridges increases (McKillop and Geeves, 1993). Second, thin filament activation affects the rate of thin filament activation and deactivation, a property referred to as “thin filament cooperativity” (Kampourakis et al., 2016). Third, myosin heads can transition from a ‘disordered-relaxed state’ or ON state in which they can bind to actin, to a ‘super-relaxed state’ or OFF state in which they cannot (Irving, 2017). The rate at which myosin heads enter and exit the super-relaxed state is force-dependent, enabling the recruitment of additional myosin heads as force increases. Cross-bridge force therefore influences thick filament activation (i.e. the fraction of cross-bridges in the disordered relaxed state), a property referred to as “thick filament cooperativity” (Campbell et al., 2018). Muscle shortening results in a reduction in both the number of attached cross-bridges and the force per cross-bridge, and thereby affects both thin and thick filament activation. Thin and thick filament cooperativity may thereby contribute to muscle thixotropy. To date, it remains unclear whether biophysical models require cooperative myofilament dynamics to reproduce muscle thixotropy, because previous models either did not include such dynamics (Campbell and Moss, 2000; Campbell and Moss, 2002), or were not tested for predicting short-range stiffness across a broad range of conditions including activation, stretch amplitudes and time histories (Blum et al., 2020; Campbell, 2014; Campbell et al., 2018; Lakie and Campbell, 2019; Liu et al., 2024; Simha and Ting, 2024). Therefore, we assessed the need for cooperative myofilament activation to capture short-range stiffness and its history dependence across a range of behaviorally relevant conditions.

**Biophysical cross-bridge models may require an additional forcibly-detached cross-bridge state to explain short-range stiffness.** Early cross-bridge models (Huxley, 1957; Zahalak, 1981) captured steady-state force-velocity curves, while only simulating two cross-bridge states: attached or detached. However, these simple models poorly captured force transients during stretch (Cole et al., 1996; van den Bogert et al., 1998). More specifically, these models predicted a large increase in force during stretch followed by a decay, while empirical forces rise during stretch and plateau with further lengthening (Cole et al., 1996; van den Bogert et al., 1998). More complex biophysical models sometimes include a ‘forcibly-detached’ cross-bridge state (Lombardi and Piazzesi, 1990), which may be required for reproducing forces during both shortening and lengthening. A forcibly-detached stated is thought to be associated with a faster re-attachment rate, required to reproduce the plateau phase of the force response. This state does not influence forces during shortening, as it can only be entered when cross-bridges are stretched beyond a ‘critical stretch’. However,, it remains unclear whether the inclusion of a forcibly detached cross-bridge state is required to explain short-range stiffness and its history dependence. Therefore, we explored the effect of including a forcibly-detached cross-bridge state on the ability to capture short-range stiffness.

**Approximating the cross-bridge distribution with a Gaussian reduces computational cost, but its effects on the ability to capture short-range stiffness and muscle thixotropy remain unclear.** Existing biophysical muscle models vary in the way they describe the strain distribution of attached cross-bridges. Some solution methods discretize the distribution into a finite number of bins (Campbell, 2014; van Soest et al., 2019), while others approximate the distribution using an analytical function such as a Gaussian (Zahalak, 1981; van der Zee et al., 2024). Until now, the Gaussian approximation has only been used in two-state models without cooperative myofilament activation. It remains unclear whether a Gaussian approximation also yields sufficiently accurate simulations for more complex models. Considering that numerical complexity is an important consideration in musculoskeletal simulations (Falisse et al., 2019), a better understanding of how solution methods affect the accuracy of simulations is needed before biophysical models are implemented in movement simulation. Therefore, we explored the effect of approximating the cross-bridge distribution with a Gaussian on the ability to capture short-range across conditions.

**Here, the aim was to identify a muscle model suitable for musculoskeletal simulations of unsteady and perturbed movements based on its ability to capture muscles’ history-dependent resistance to stretch.** To this end, we evaluated muscle force predictions from various muscle models against existing data on muscle forces for a broad range of behaviorally relevant movement histories and activation levels, recently documented in rat soleus muscle fibers (Horslen et al., 2023). We assessed the effects of including cooperative myofilament activation, a forcibly detached cross-bridge state and a Gaussian approximation of the cross-bridge distribution on the ability to capture short-range stiffness and its history dependence. We also tested a Hill-type model for reference. We found that biophysical models with cooperative myofilament activation and a forcibly-detached cross-bridge state yielded best agreement with experimental data. Approximating the cross-bridge distribution with a Gaussian had little effect on the force transients but greatly reduced computational time.

## Methods

We tested three different biophysical muscle models of increasing complexity against *in vitro* data from permeabilized rat soleus muscle fibers for a range of behaviorally relevant stretches. Data included force and length trajectories of 11 permeabilized rat soleus muscle fibers for stretch-shortening-stretch protocols at 4-6 different calcium concentrations, 4-7 different amplitudes of the first stretch, and 5-7 isometric recovery times before the second stretch, for a total of 80-196 conditions per fiber. Three biophysical models were included: (1) two cross-bridge states (i.e. attached, detached) without cooperative myofilament activation, (2) two cross-bridge states with cooperative activation of both thin and thick filaments, and (3) three cross-bridge states (i.e. attached, detached and forcibly detached) with cooperative activation of both thin and thick filaments. We evaluated all models both with and without a Gaussian approximation of the cross-bridge distribution. Model parameters were fit on a subset of data (two stretch-shortening-stretch protocols across all activation levels) and evaluated on the remaining data. We first describe the experimental data in more detail and then describe the biophysical muscle models.

### Experimental data

We fitted and evaluated biophysical muscle models on existing data from a broad range of stretch-shortening-stretch trials on 11 permeabilized rat soleus muscle fibers (Horslen et al., 2023). This data included isometric trials and stretch-shortening-stretch ramps at various calcium concentrations. Empirical forces were normalized with respect to the maximal isometric force F_0_, obtained during the isometric trials at pCa = 4.5. The stretch-shortening-stretch trials featured an isometric phase, followed by an isokinetic conditioning stretch, an isokinetic shortening phase back to the pre-conditioning length, an isometric recovery phase and an isokinetic test stretch followed by another isometric phase (Fig. 2). Conditions differed in their calcium concentration (Activation, Fig. 2A), conditioning stretch-shortening amplitude (AMP, Fig. 2B), and isometric recovery time (RT, Fig. 2C). Most fibers (n = 8) were tested at 4 AMPs and 5 RTs at 5-6 pCas. A subset of fibers (n = 3) was tested at 7 AMPs and 7 RTs for 4-5 pCas (Fig. 2D-E). Some AMP-RT combinations were excluded to avoid fatigue. All fibers also had a reference condition, without a conditioning stretch. The reader is referred to the original publication (Horslen et al., 2023) for more details on experimental data. We selected two stretch-shortening conditions across activation levels for fitting model parameters (“Fitting”, Fig. 2D), one with a large conditioning stretch amplitude and short recovery time resulting in large short-range stiffness reductions (AMP = 3.83% L_0_, RT = 0.1 s) and the other without conditioning stretch (AMP = 0% L_0_, i.e. no conditioning stretch). Whereas the first stretch is identical in both conditions, it is followed by shortening in the case of a conditioning stretch and by an isometric contraction in the absence of a conditioning stretch. The remaining trials were used to validate model predictions (“Validate”, Fig. 2D).

**Figure 2.**
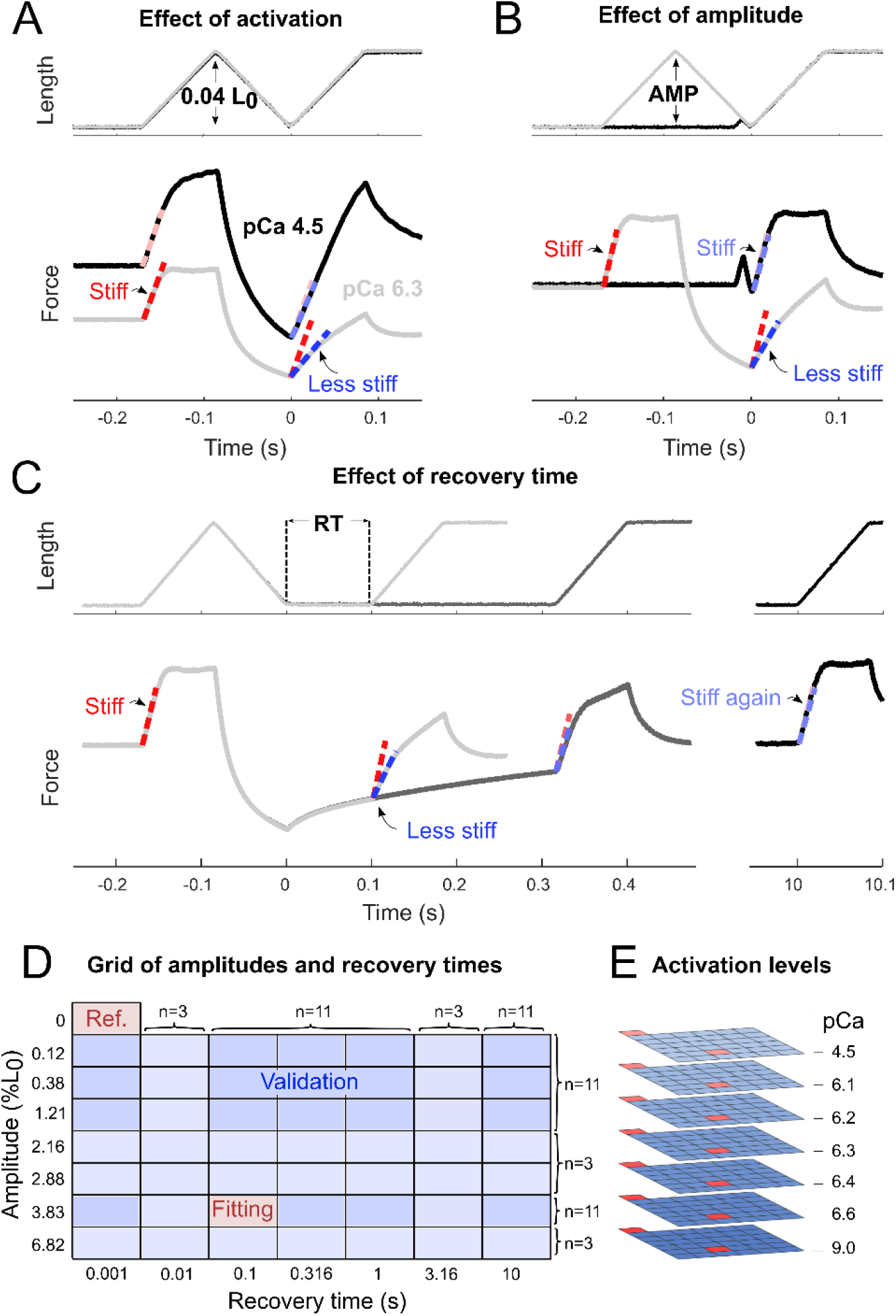
Experimental conditions. Data were obtained using stretch-shortening-stretch protocols that differed in conditioning stretch amplitude, recovery time, and calcium concentration. **A:** Muscle fiber length and force trajectories for protocols at constant stretch amplitude and recovery time but variable activation. **B:** Muscle fiber length and force trajectories for protocols at constant activation and recovery time but variable stretch amplitude (AMP). **C:** Muscle fiber length and force trajectories for protocols at constant amplitude and activation but variable recovery time (RT). **D:** Grid of tested amplitudes and recovery times. Models were fitted on data from two trials per calcium concentration (red) and evaluated on data from the remaining trials (blue). Most fibers (n = 8) had grids with 4 amplitudes and 5 recovery times. A subset of fibers (n = 3) had grids with 7 amplitudes and 7 recovery times. Empirical and modeled short-range stiffness during stretch-shortening-stretch protocols was expressed relative to short-range stiffness during a reference stretch trial without prior stretch-shortening (Ref). **E:** Grids of amplitudes and recovery times were repeated for up to 7 different calcium concentrations.

### Data processing

Data was analyzed and processed using custom MATLAB-based software (MathWorks, Natick, MA, USA). Empirical forces during stretch-shortening-stretch trials were scaled such that the initial isometric force equaled that during the corresponding pCa-matched isometric trial to remove potential fatigue effects. Short-range stiffness was quantified as the average slope of the force-length plot during the first 10 ms after stretch onset. The average slope was determined by fitting a straight line (MATLAB’s *polyfit*) to the force versus length data. Muscle thixotropy was quantified using the relative short-range stiffness, defined as the ratio between the short-range stiffness of the test stretch compared to that of an unconditioned stretch. Relative short-range stiffnesses of 0 and 1 respectively indicate maximal and minimal muscle thixotropy. For each fiber, relative stiffness estimates at different calcium concentrations were assigned to one of the following categories: Passive (< 0.07 F_0_), Low activation (0.07 – 0.25 F_0_), Intermediate activation (0.25 – 0.7 F_0_) and High activation (> 0.7 F_0_).

### Biophysical dynamics

#### Biophysical models

We explored a range of biophysical muscle models that describe the mechanisms that are thought to underly muscle short-range stiffness and its history dependence. Considering the potential role of cooperative myofilament dynamics in muscle thixotropy, we tested a previously described model that includes these dynamics (Campbell et al., 2018), referred to as **XB coop model**. We also tested a model without such dynamics (Lemaire et al., 2016), referred to as **XB model**. We also evaluated the effects of adding a forcibly detached cross-bridge state, referred to as **XB coop + FD model**. Lastly, we evaluated all models with both a discretized and an approximated cross-bridge distribution solution method.

#### Cross-bridge cycling

All biophysical muscle models considered here are based on the cross-bridge model of Huxley (1957), which tracks the fraction *n* and the strain *x* of attached cross-bridges. This model originally only considered muscle fibers at maximal activation but has been updated over the years to incorporate submaximal activation (Zahalak, 1981). Here, we adopt the following version of the state equation (Campbell et al., 2018):

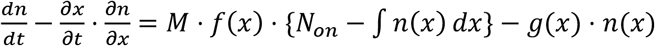

Here, *n*(*x*) is the fraction of attached cross-bridges *n* for each strain *x, M* is the fraction of myosin heads in the disordered-relaxed state (see ‘Thick filament activation’, below), *N*_*on*_ is the fraction of thin filament binding sites in the active state (see ‘Thin filament activation’, below), and the integral ∫ *n* (*x*)*dx* indicates the fraction of attached cross-bridges. Functions *f*(*x*) and *g*(*x*) are the strain-dependent attachment and detachment rate functions:

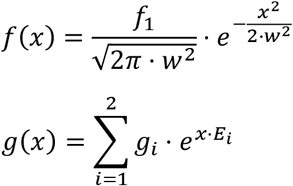

Here, attachment function *f*(*x*) is a Gaussian that is specified by three parameters: mean *ϵ*_*ps*_, standard deviation *w* and amplitude *f*_1_. A Gaussian formulation was chosen because it reflect the Brownian motion of the myosin head before attachment (Campbell et al., 2018). Detachment function *g*(*x*) is the sum of two exponentials that are each scaled by a rate constant (*g*_1_ and *g*_2_) and shaped by a dimensionless exponential strain dependence (*E*_1_ and *E*_2_). An exponential formulation for the detachment function was chosen in agreement with empirical estimates of cross-bridge detachment and the strain-dependent release of ADP (Liu et al., 2024). Cross-bridge strain *x* is normalized and centered relative to the size of the cross-bridge power stroke *ϵ*_*ps*_:

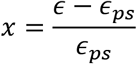

Here, *ϵ* and *x* are the absolute cross-bridge elongation and the normalized cross-bridge strains, respectively. Because of centering, normalized cross-bridge force is proportional to the sum of the zero-order and first-order moments of the cross-bridge distribution *n*(*x*):

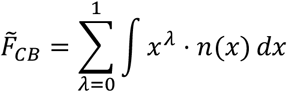

Here, 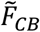 is the cross-bridge force, normalized with respect to the force obtained when all cross-bridges are attached with a strain equal to that of the power stroke (i.e. *ϵ* = *ϵ*_*ps*_, and *x* = 0). Optionally, two separate pathways were included by formulating forcible detachment and reattachment functions *ϕ*(*x*) and Γ(*x*):

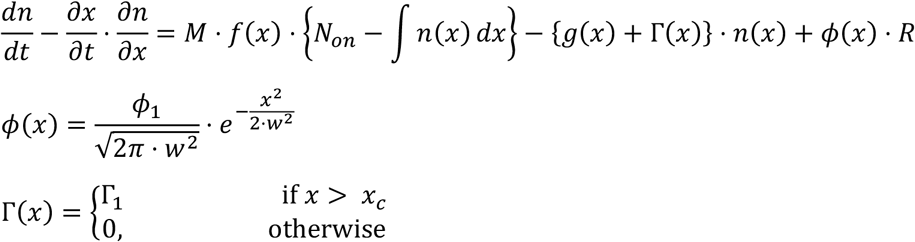

Here, *R* is the fraction of cross-bridges in the forcibly detached state. Its dynamics are described by the following equation:

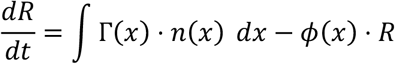

#### Thin filament activation

Thin filament dynamics are implemented using the following state equation (Campbell et al., 2018):

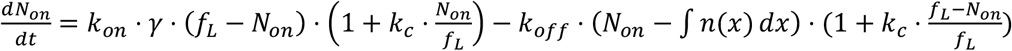

Here, *γ* is the calcium concentration, and *k*_*on*_ and *k*_*off*_ are activation and deactivation rate constants, respectively. Parameter *k*_*c*_ is a constant that defines the strength of thin filament cooperativity. If *k*_*c*_ is greater than zero, active binding sites induce other sites to activate, whereas inactive sites accelerate deactivation. The term *N*_*on*_ - ∫ *n* (*x*)*dx* results in cross-bridge-mediated thin filament activation. More specifically, thin filament binding sites can only switch off if they are not occupied by cross-bridges. In the model without cooperative myofilament activation, *N*_*on*_ depends on calcium concentration *γ* via a sigmoidal saturation function (Curtin et al., 1998):

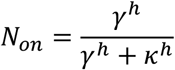

Here, κ sets the horizontal position of the sigmoid, and *h* is the Hill coefficient that sets the steepness.

#### Thick filament activation

Thick filament dynamics are implemented using the following state equation (Campbell et al., 2018):

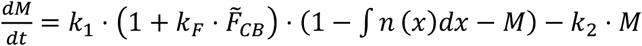

Here, *k*_1_ and *k*_2_ are forward and backward rate constants, respectively. Parameter *k*_*F*_ specifies the sensitivity of the forward rate to normalized cross-bridge force ^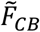^. In the model without cooperative myofilament activation, cross-bridges are not allowed to leave the disordered-relaxed state and enter the super-relaxed state. This is implemented by setting *k*_2_ = 0, and setting the initial value of *M* equal to 1.

#### Interface with elastic elements

Biophysical muscle dynamics are interfaced with parallel and elastic tissues (Fig. 3B), similar to previously described (Lemaire et al., 2016; van der Zee et al., 2024). However, unlike the most common Hill-type configuration, we chose a configuration in which the parallel element is also in parallel with the series element. This choice is inspired by the corresponding biological configuration (inset, Fig. 3B), in which titin and myosin are (at least partially) in parallel with actin. Cross-bridge force *F*_*CB*_ is combined with forces from elastic tissues to yield an overall muscle fiber force *F*. The parallel elastic element is a linear spring, specified by two parameters (i.e. stiffness *k*_*p*_ and resting length *L*_*p*,0_). The series elastic element is an exponential spring, specified by two parameters (i.e. coefficient *σ*, exponent *L*_*c*_).

**Figure 3.**
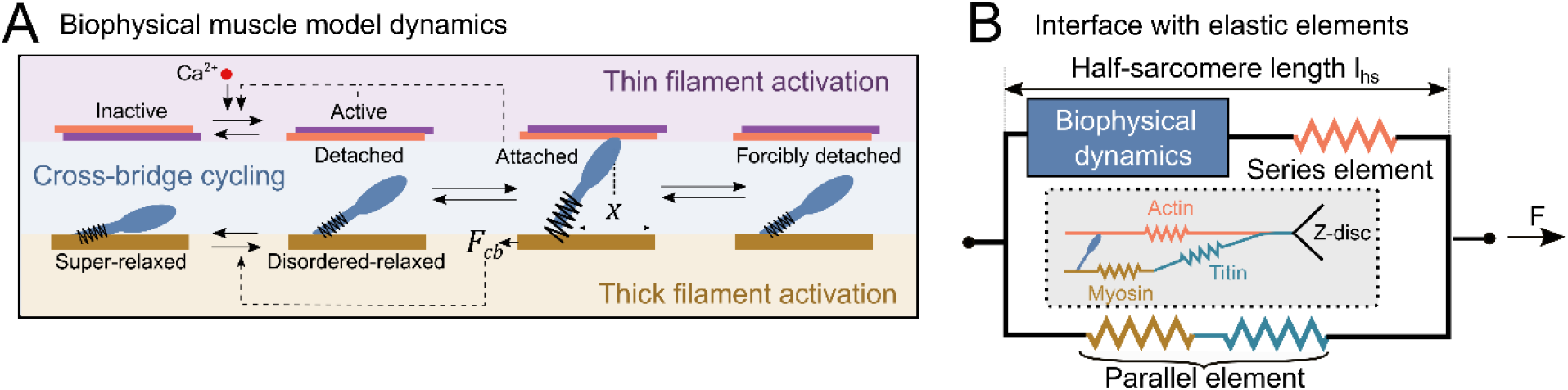
Biophysical muscle model dynamics and interface with elastic elements. **A**. Biophysical muscle models distinguish between dynamics for thin filament activation, thick filament activation, and cross-bridge cycling. Thin filament activation depends on myoplasmic calcium [Ca]^2+^, and is optionally mediated by the fraction of active thin filament binding sites and the fraction of attached cross-bridges (dashed lines). Thick filament activation is mediated by cross-bridge force *F*_*CB*_ (dashed lines), which depends on cross-bridge strain *x*.. **B**. Biophysical muscle model dynamics are interfaced with series and parallel elastic elements. The inset shows the biological structures corresponding to the modelled elastic elements.

#### Gaussian approximation

Cross-bridge cycling dynamics are resolved either using a distribution-moment approximation, or a discretized cross-bridge distribution solution method. For the former, a Gaussian approximation was applied to the distribution of attached cross-bridges *n*(*x*), yielding:

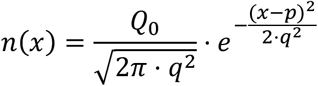

Here, *Q*_0_ is the zero-order moment of the distribution, while *p* and *q* are the mean and standard deviation, respectively. *p* and *q* can be calculated from the moments:

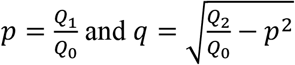

This approximation yields a set of three state derivative equations:

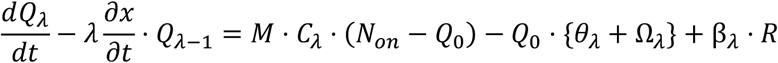

Here, *Q*_*λ*_, *C*_*λ*_, *θ*_*λ*_, Ω_*λ*_ and β_*λ*_ are the *λ*-order moments of the cross-bridge distribution *n*(*x*), the attachment function *f*(*x*), and a zero-order normalized cross-bridge distribution multiplied by the detachment function *g*(*x*), zero-order normalized cross-bridge distribution multiplied by the detachment function Γ(*x*), and the attachment function *ϕ*(*x*), respectively. The 0^th^ through 2^nd^-order moments are considered, yielding a gaussian approximation and three models states: *Q*_0_, *Q*_1_ and *Q*_2_. *Q*_−1_ is set equal to 0. For the rate functions used here, the moments *C*_*λ*_, *θ*_*λ*_, Ω_*λ*_ and β_*λ*_ have analytical solutions (see Appendix). Strain rate 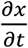 can be computed from the model states, as described previously (van der Zee et al., 2024). For discretized cross-bridge models, cross-bridge populations were evaluated with a 0.71 nm resolution over a strain range of ±15 *ϵ*_*ps*_, yielding 500 strain bins.

### Model parameters

Model parameters fall into one of the following categories: (1) cross-bridge cycling, (2) thin and thick filament activation, (3) elasticity. We next describe the parameters within each category in further detail.

#### Cross-bridge cycling

The rates of cross-bridge attachment and detachment are specified by strain-dependent functions *f*(*x*) and *g*(*x*), and optionally *ϕ*(*x*) and Γ(*x*). The mean of the attachment function *f*(*x*) corresponds to the size of the cross-bridge power stroke (*ϵ*_*ps*_), estimated at 10 nm (Finer et al., 1994). The standard deviation *w* of the attachment function *f*(*x*) reflects random Brownian motion when the myosin head binds to actin, which is estimated at 1 nm based on the experimental temperature (22 C) and a crossbridge stiffness of 0.5 pN/nm (Brenner et al., 2012; Percario et al., 2018). This yields a cross-bridge power-stroke force of 5 pN and a maximum work output of 50 zJ, both in agreement with previous estimates (Barclay, 2015). The strain dependence of the detachment function *g*(*x*) at negative strains reflects the force dependence of the ADP release rate, which is assumed to have a dimensionless coefficient *E*_1_ of 2 (Liu et al., 2024). The strain dependence of the detachment function *g*(*x*) at positive strains reflects cross-bridge ripping (Liu et al., 2024), with a dimensionless coefficient *E*_2_. In case forcible detachment is included, another detachment pathway is added through detachment function Γ(*x*). A step function is used, specified by a fast rate constant *r* (set to 3000 s^-1^) for strains beyond a critical stretch *ϵ*_*c*_. Subsequent attachment occurs through the attachment function *ϕ*(*x*), a gaussian that is assumed to have the same standard deviation *w* as *f*(*x*), but a different mean *ϵ*_*FD*_ and a higher amplitude *ϕ*_1_ (set at 1000 s^-1^). The following cross-bridge parameters were fitted on experimental data from the fitting trials for all XB models: *f*_1_, *g*_1_, *g*_2_, and *E*_2_. For the XB coop + FD model, two additional parameters were fitted: *ϵ*_*c*_ and *ϵ*_*FD*_.

#### Thin and thick filament activation

The rates of thin and thick filament activation are specified by two state equations. XB coop and XB coop + FD models have cooperative thin filament activation, which depends on three parameters: forward rate *k*_*on*_, backward rate *k*_*off*_ and cooperative parameter *k*_*c*_. The backward rate for thin filament activation *k*_off_ was estimate at 80 s^-1^, based on temperature corrected estimates of the calcium-troponin dissociation rate in mouse soleus muscle fibers (Baylor and Hollingworth, 2003). Thick filament activation is specified by three parameters: forward rate *k*_1_, backward rate *k*_2_ and force-dependent parameter *k*_*F*_. The forward and backward rates *k*_1_ and *k*_2_ were set to 6.17 s^-1^ and 200 s^-1^, based on previous estimates (Campbell et al., 2018). For the models with cooperative myofilament activation, the following myofilament activation parameters were fitted on experimental data from the fitting trials: *k*_*on*_, *k*_*c*_ and *k*_*F*_. The Hill-type and XB model do not have these cooperative dynamics but instead rely on a sigmoidal activation function specified by parameters *h* (steepness) and κ (horizontal shift) to translate calcium concentration into muscle activation. For these models, parameters *h* and κ were fitted on experimental data from the fitting trials.

#### Elasticity

The parallel elastic element resting length *L*_*p*,0_ and stiffness *k*_*p*_ were fit on passive ramp stretches at pCa = 9.0 for each individual fiber. *L*_*p*,0_ and *k*_*p*_ differed between fibers, but not between models. Series elasticity parameters σ and *L*_*c*_ were fitted on experimental data from the fitting trials for each model.

#### Hill-type model

As a benchmark, we also tested a standard Hill-type model. The same model configuration was used (Fig. 3B) in combination with a force-velocity relation to capture the dependence of force on stretch velocity (instead of biophysical dynamics). Default force-velocity shape parameters were used (De Groote et al., 2016). Unloaded shortening velocity *v*_*max*_ was fitted on experimental data from the fitting trials.

### Parameter estimation

All unknown parameters were estimated by fitting the simulated forces to the empirical forces for to stretch-shortening-stretch conditions (“Fitting”, Fig. 3). Data from all calcium concentrations available for these two conditions was used for parameter estimation. Given the imposed length changes and calcium concentrations, the parameter values of each muscle model were estimated by minimizing a cost function composed of the sum-squared differences between the simulated forces and empirical forces. The time interval considered for fitting ranged from 0.0843 s before the onset of the conditioning stretch until 0.0843 s after the end of the test stretch. For biophysical models, parameter estimation was performed using the Gaussian approximated solution method. Optimization problems were solved using direct collocation (CasADi; Andersson et al., 2019). The resulting nonlinear optimization problems were solved using IPOPT (Wächter and Biegler, 2006). Four (out of 11) fibers were excluded from fitting because they did not include stretch-shortening-stretch trials at Intermediate and/or Maximal activations, which we found to be key to fitting. Thus, fitting was performed on data from 2 fitting trials for 7 muscle fibers. We used the estimated parameter in combination with both solution methods to evaluate forces for all trials. All code pertaining to the muscle models is available on GitHub (https://github.com/timvanderzee/biophysical-muscle-model).

### Model evaluation and predictions

The ability of muscle models to capture the empirical data was assessed in two different ways. First, the root-mean-square difference (RMSD) between modelled forces and empirical forces was computed for all experimental trials for each fiber included in the fitting set (n = 7). The RMSDs were expressed as a percentage of the maximal isometric force *F*_0_ measured at pCa = 4.5. We predicted that the RMSD would be higher for the Hill-type model than for all biophysical models. Furthermore, we predicted that biophysical models with cooperative myofilament activation would have a lower RMSD than those without cooperative myofilament activation. We also tested the effect of including the forcibly detached state and approximating the cross-bridge distribution with a Gaussian. Second, model predictions of muscle thixotropy were evaluated. To this end, we computed the deviations between the modelled (n = 7) and measured (n = 11) relative short-range stiffness across all experimental conditions. From these deviations, we computed the R^2^ to evaluate the goodness of fit of each model. To account for the different numbers of parameters in the different models, we also computed the Akaike information criterion (AIC) score of each model. The AIC score of biophysical models was expressed relative to that of the Hill-type model (ΔAIC). In a portion of the results, we separately analyzed trials with small versus large history dependence. Based on the experimental results (Horslen et al., 2023), trials with large history dependence were defined as those with Low or Intermediate activation, stretch amplitude greater than 3% L_0_ and recovery time of less than 0.5 s. Trials with small history dependence were defined as the remaining trials. We anticipated that differences between models would be more pronounced for trials with greater history dependence.

## Results

After fitting to force trajectories during two stretch conditions at different calcium levels, biophysical models captured forces across stretch-shortening conditions better than a Hill-type model, but only if cooperative myofilament dynamics were included. Neither a biophysical model lacking cooperative dynamics nor a Hill-type model captured the reduced muscle stiffness after a stretch-shorten cycle, i.e. muscle thixotropy. Adding cooperative myofilament dynamics reduced the rate of force development and the short-range stiffness after stretch-shorten cycles, and was robust across activation levels, recovery times, and pre-stretch amplitudes. Finally, adding a forcibly detached state slightly exaggerated the thixotropic behavior of the model. The cross-bridge models including cooperative dynamics could robustly capture the effects of multiple activations, stretch-shorten amplitude, and recovery times on the time course of muscle force observed experimentally in permeabilized, activated muscle fibers. Below, we first describe qualitative differences in force trajectories simulated based on different models. Then, we quantitatively evaluated how well different models capture force trajectories and muscle thixotropy across conditions.

**Cross-bridge dynamics alone were insufficient to explain muscle forces during stretch-shortening at submaximal activation.** Whereas both the XB model (green lines, Fig. 4) and Hill-type model (red lines, Fig. 4) captured forces during the fitting conditions at maximal activation relatively well (Fig. 4A), they failed to capture the reduction in force after a preconditioning stretch at submaximal activation (Fig. 4B). At submaximal activation, both these models underestimated forces during the conditioning stretch and overestimated forces during the test stretch (Fig. 4B), which might be a trade-off resulting from the inability to capture the reduction in force due to a conditioning stretch. Observations were similar for testing conditions (Fig. 5, exemplar conditions with shorter/longer recovery times, green and red traces). Overall, both the XB model and the Hill-type model did not capture muscle forces during stretches at submaximal activation.

**Figure 4.**
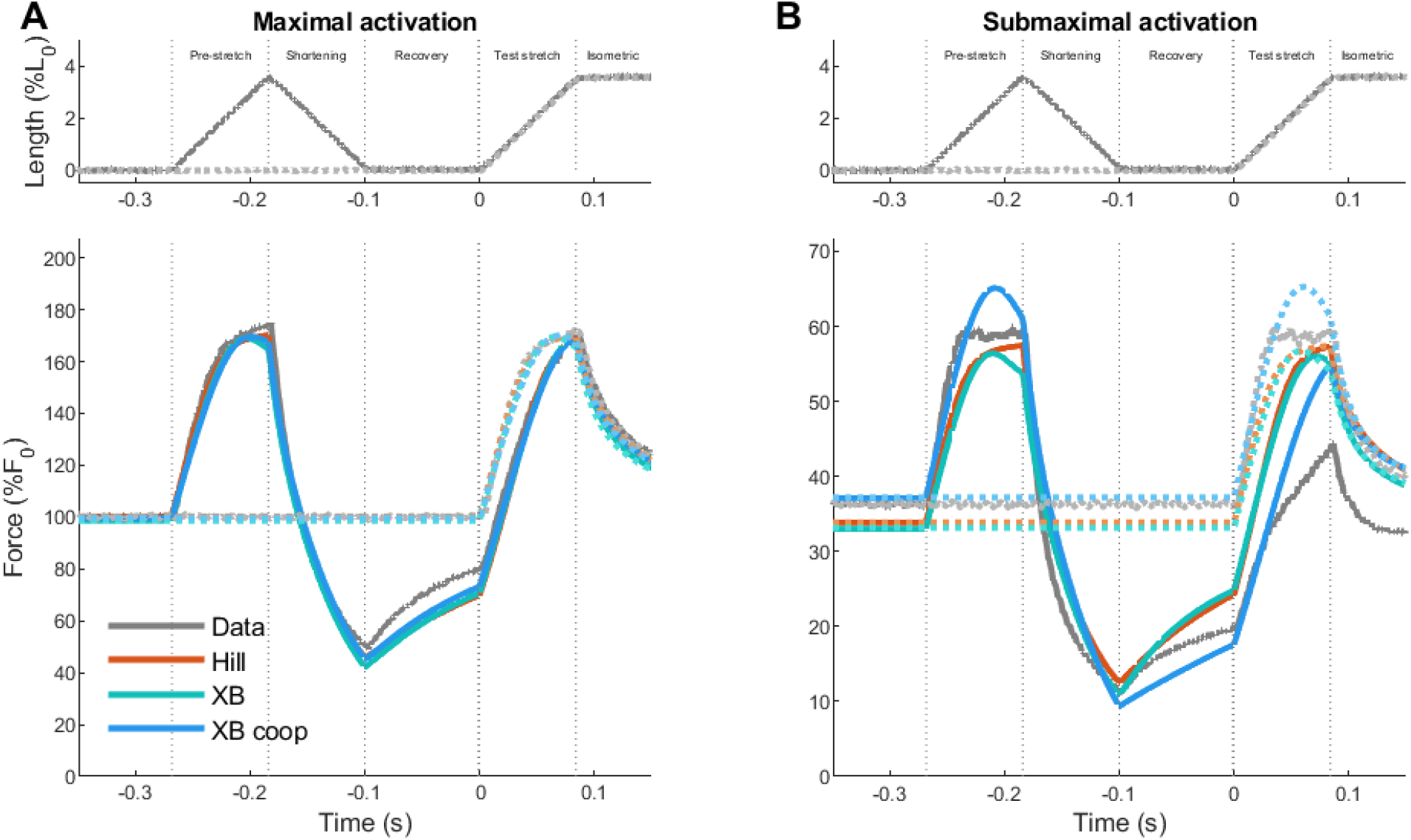
Modelled and measured forces for fitting trials: effect of cooperative dynamics. Top row shows measured fiber length changes for a condition with a pre-stretch (darker solid lines) and without a pre-stretch (lighter dotted lines). Bottom row shows the corresponding empirical forces (“Data”, grey), alongside forces of Hill-type model (“Hill”, red), cross-bridge model (“XB”, green), cross-bridge model with cooperative dynamics (“XB coop”, blue). **A**: maximal activation (pCa 4.5). **B**: submaximal activation (pCa 6.3).

**Figure 5.**
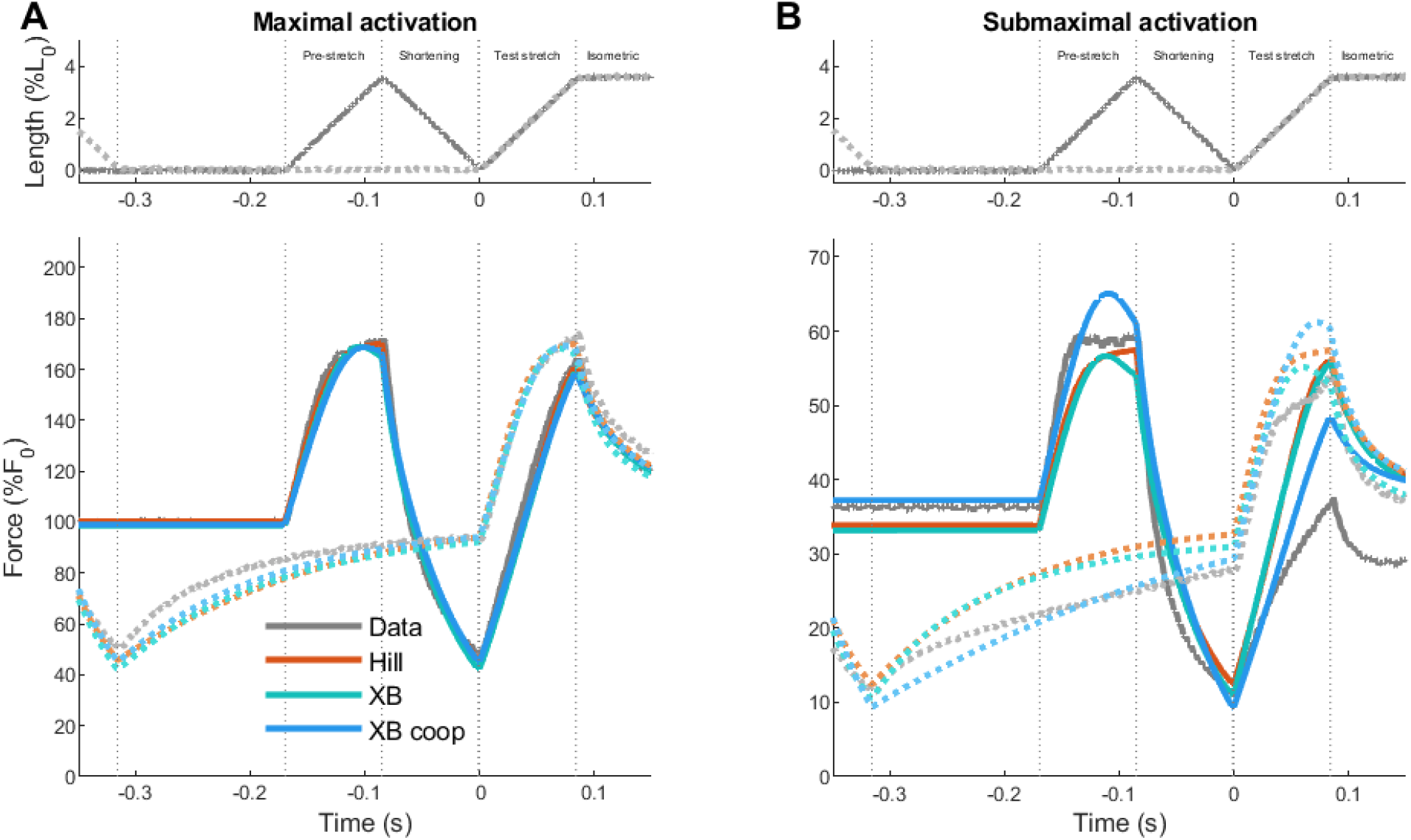
Modelled and measured forces for two testing trials: effect of cooperative dynamics. Top row shows measured fiber length changes for a condition with short recovery time (darker solid lines) and long recovery time (lighter dotted lines). Bottom row shows the corresponding empirical forces (“Data”, grey), alongside forces of Hill-type model (“Hill”, red), cross-bridge model (“XB”, green), cross-bridge model with cooperative dynamics (“XB coop”, blue). **A**: maximal activation (pCa 4.5). **B**: submaximal activation (pCa 6.3).

**Adding cooperative myofilament dynamics improved predictions of muscle forces during stretch-shortening at submaximal activation.** At maximal activation, forces predicted by the XB coop model resembled those predicted by the XB model for both fitting conditions (Fig. 4A, blue versus green lines) and testing conditions (Fig. 5A, exemplar conditions with shorter/longer recovery times, blue versus green traces). However, at submaximal activation, the XB coop model better captured forces compared with both Hill-type and XB models for both fitting conditions (Fig. 4B) and testing conditions (Fig. 5B). For both the testing condition and the fitting condition with conditioning stretch at submaximal activation, the XB coop model overestimated the force during both stretches (Fig. 4B). However, unlike the Hill-type model and XB model, the XB coop model captured the slow force recovery during the isometric phase following shortening in these conditions. Furthermore, unlike the Hill-type and XB model, the XB coop model predicted a reduction in the force during the test stretch compared with the force during the conditioning stretch. Thus, cross-bridge dynamics and cooperative myofilament dynamics are required for accurately predicting history-dependent muscle forces during stretches at submaximal activation.

**Combining cooperative myofilament dynamics with a forcibly detached cross-bridge yielded greater history dependence at submaximal activation.** At maximal activation, the XB coop + FD model predicted slightly steeper increases in force during both the conditioning and test stretches compared with the XB coop model, which better match experimental increases (Fig. 6A, exemplar test conditions at maximal activation). Both models differed most in the predictions of force during the plateau phase of the first (unconditioned) stretch after the initial steep increase in force. Surprisingly, the XB coop model captured the plateau better than the XB coop + FD model. At submaximal activation, the XB coop + FD model predicted a similar increase in force during pre-stretch as the XB coop model, but a slightly slower force increase during the test stretch (Fig. 6B). These predictions imply greater history dependence that was more in line with experimental data. Thus, XB coop + FD model yielded slightly more history dependence but did not yield better force predictions.

**Figure 6.**
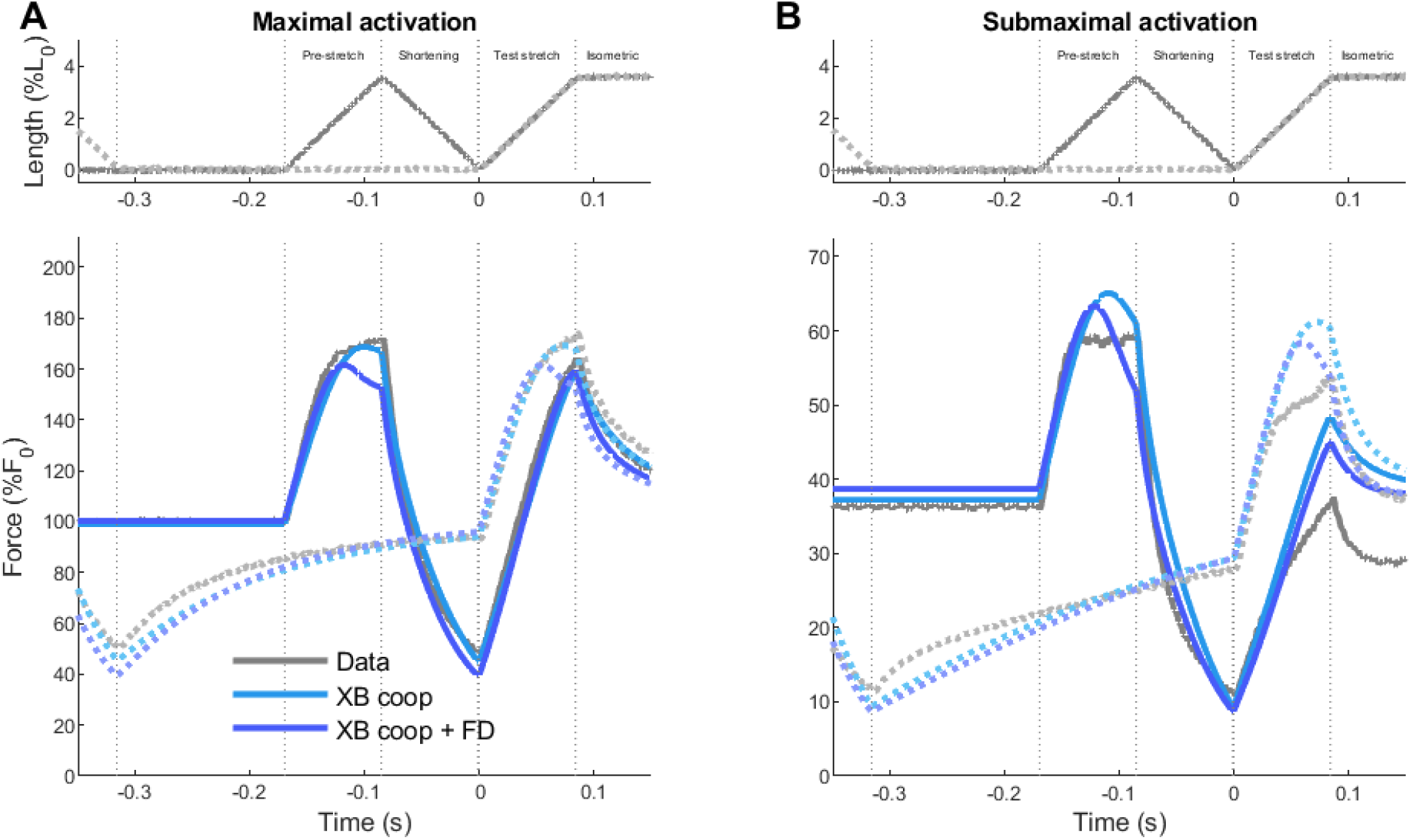
Modelled and measured forces for two testing trials: effect of forcibly detached state. Top row shows measured fiber length changes for a condition with short recovery time (darker solid lines) and long recovery time (lighter dotted lines). Bottom row shows the corresponding empirical forces (“Data”, grey), alongside forces of Hill-type model (“Hill”, red), cross-bridge model (“XB”, green), cross-bridge model with cooperative dynamics (“XB coop”, light blue), and cross-bridge model with cooperative dynamics and a forcibly detached state (“XB coop + FD”, dark blue). **A**: maximal activation (pCa 4.5). **B**: submaximal activation (pCa 6.3).

**Cooperative myofilament dynamics is key to accurately predicting forces during stretch-shortening, whereas additionally modeling a forcibly detached state has little effect on the overall agreement between model and experimental forces.** We computed RMSDs between model forces and experimental forces across conditions, averaged over all fibers for which we had sufficient data to estimate parameters (n = 7, Fig. 7). In line with exemplar force trajectories, predictions from the XB coop model (light blue triangles) and XB coop + FD model (dark blue circles) agreed better with data than predictions from the XB model (green hexagons) and the Hill-type model (red squares). Differences were apparent both for conditions with large short-range stiffness reduction (closed symbols) and for conditions with small short-range stiffness reduction (open symbols). Differences in RMSD between models were more pronounced during the conditioning and test stretch, than averaged over the entire force trajectory. Errors were very similar for the Hill and XB models, and for the XB coop and XB coop + FD models. Force predictions of the XB coop and XB coop + FD models generally matched experimental forces better than force predictions from the Hill and XB models, particularly during stretching.

**Figure 7.**
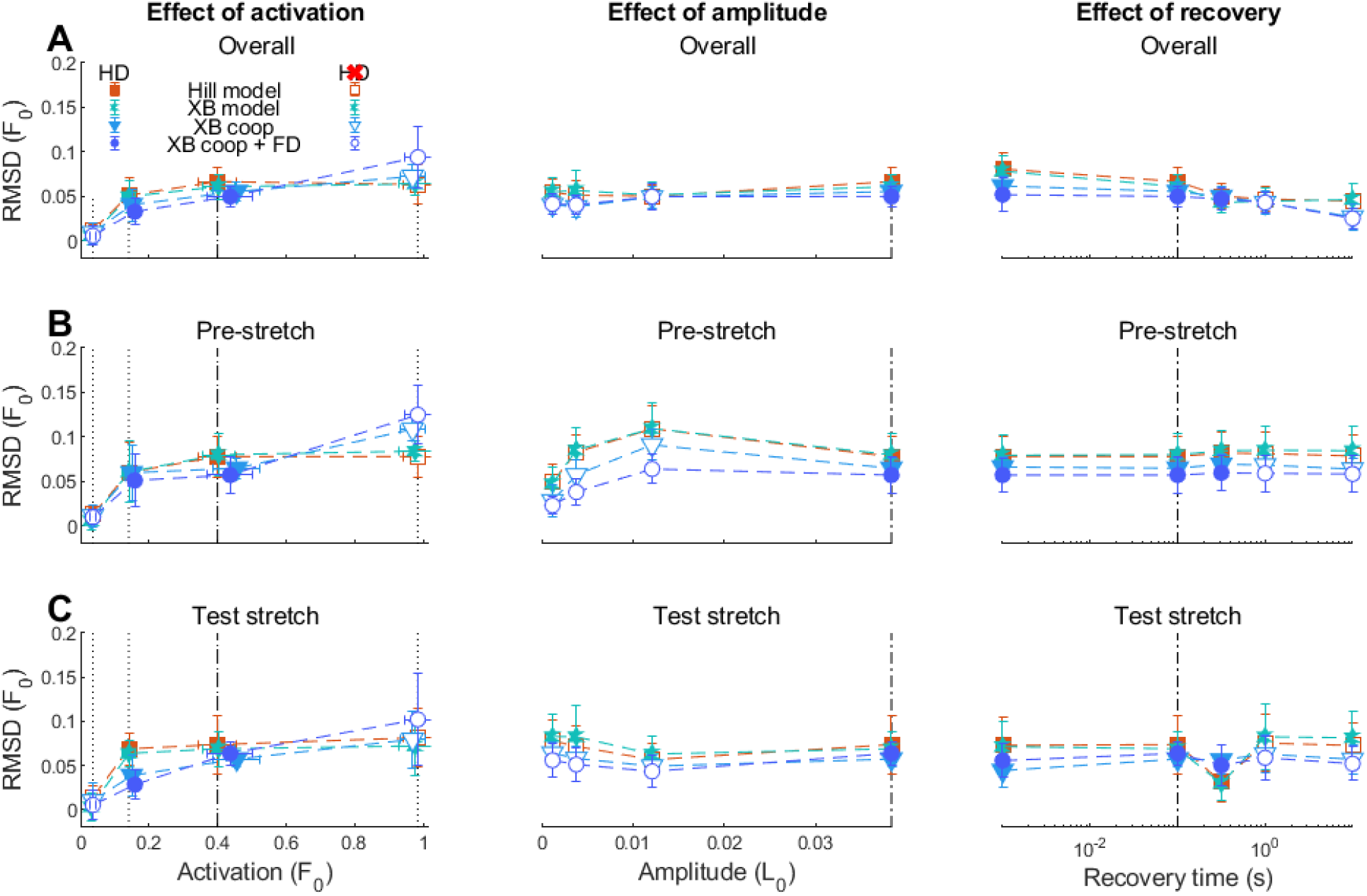
Root-mean-squared deviations between model and experimental forces. RMSDs are shown for Hill-type model (“Hill”, red), cross-bridge model (“XB”, green), cross-bridge model with cooperative dynamics (“XB coop”, blue), and cross-bridge model with both cooperative dynamics and a forcibly detached state (“XB coop + FD”, dark blue). Filled symbols indicate trials with considerable short-range stiffness reductions (i.e. history dependence), open symbols indicate trials with little to no short-range stiffness reductions (i.e. no history dependence). Left column: effect of activation at constant amplitude and recovery time. Right column: effect of amplitude at constant activation and recovery time. Right column: effect of recovery time at constant activation and amplitude. Dashed vertical lines indicate the condition common to all three sets of conditions, dotted vertical lines indicate fitting trials (dashed-dotted indicate both). **A**: Entire stretch-shortening protocol. **B**: Pre-stretch phase. **C**: Test stretch.

**Modeling cooperative myofilament activation was important to capture short-range stiffness changes across conditions.** Relative short-range stiffness predictions of the XB coop (Fig. 8, light blue lines) and XB coop + FD models (dark blue lines) closely matched experimental values, while those from the XB (green lines) and Hill-type models (red lines) did not. The XB coop and XB coop + FD models captured the effect of muscle activation (R^2^ = 0.45 – 0.86, Fig. 8A), stretch amplitude (R^2^ = 0.70 – 0.71, Fig 8B), and recovery time (R^2^ = 0.79 −0.91, Fig. 8C) on relative short-range stiffness. In contrast, the XB model did not capture any of these effects (R^2^ < 0, Fig 8A-C). The Hill-type model partially captured the effect of amplitude (R^2^ = 0.47, Fig. 8B) but did not capture the effect of activation and recovery (R^2^ < 0, Fig. 8A and Fig. 8C). Both the XB and Hill-type model incorrectly predicted a continued decline in relative short-range stiffness instead of the experimental U-shaped relation between relative short-range stiffness and muscle activation (red and green lines, Fig. 8A). The Hill-type model correctly predicted that relative short-range stiffness decreases with larger amplitude (Fig. 8B) and shorter recovery (Fig. 8C) but underestimated the magnitude of these effects. The XB model did not predict a decrease in short-range stiffness with greater amplitude or shorter recovery. Thus, incorporating cooperative myofilament dynamics improves the ability to capture short-range stiffness across a broad range of stretch-shorten conditions.

**Figure 8.**
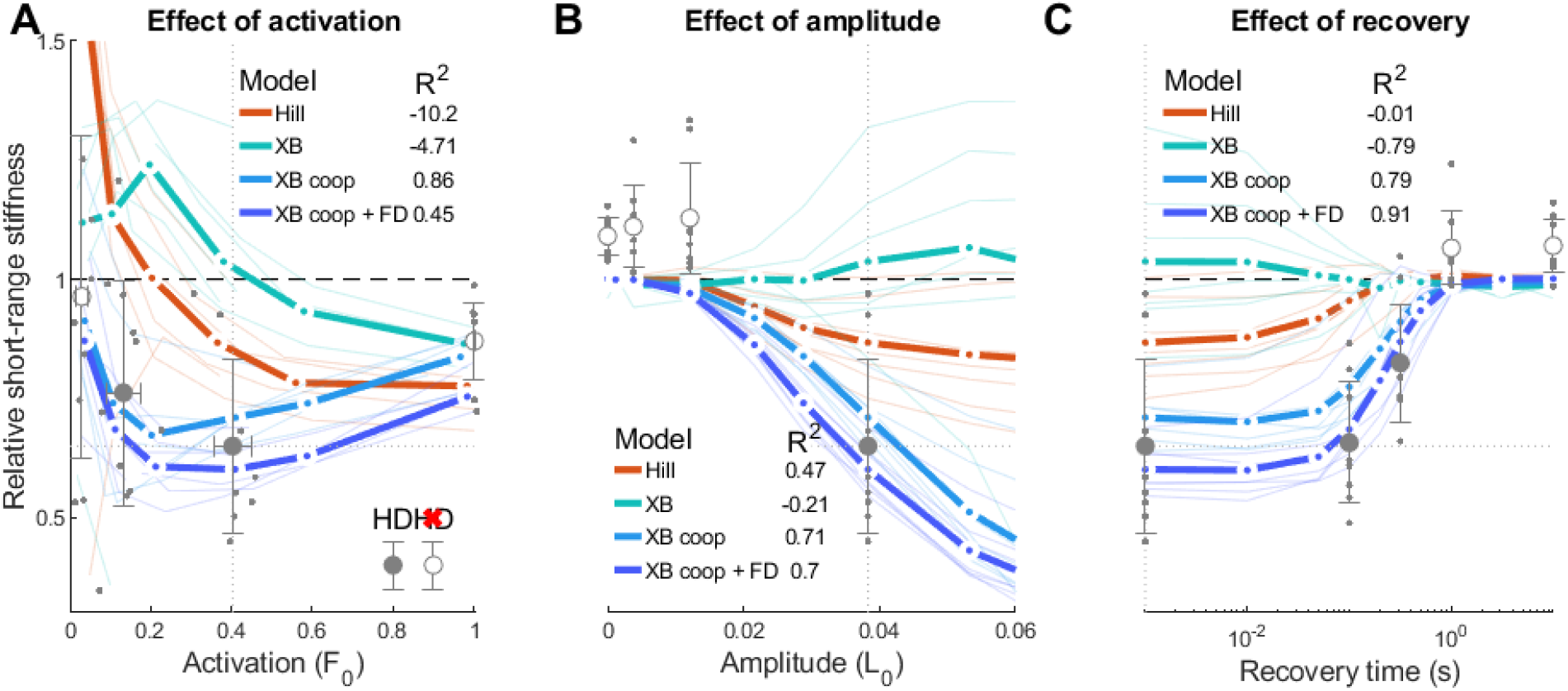
Empirical and model relative short-range stiffness across muscle activations, stretch amplitudes and recovery times. Dotted lines indicate the condition common to all three sets of conditions. Model predictions are shown for Hill-type model (“Hill”, red), cross-bridge model (“XB”, green), cross-bridge model with cooperative dynamics (“XB coop”, blue), and cross-bridge model with both cooperative dynamics and a forcibly detached state (“XB coop + FD”, dark blue). Model predictions are shown for fibers in the fitting set (n = 7), both averaged across these fibers (thick lines) and for individual fibers (thin lines). Data is shown both averaged across all fibers (n = 11, grey circles), and for individual fibers (grey dots). Filled circles indicate trials with considerable short-range stiffness reductions (i.e. history dependence, HD), open symbols indicate trials with little to no short-range stiffness reductions (i.e. no history dependence, no HD). **A**. Effect of activation at constant amplitude and recovery time. **B**. Effect of amplitude at constant activation and recovery time. **C**. Effect of recovery time at constant activation and amplitude.

**A biophysical muscle model with cooperative myofilament activation and a forcibly detached state predicts muscle thixotropy across behaviorally relevant conditions.** At submaximal activation, biophysical model with cooperative myofilament activation reproduced the observation that relative short-range stiffness (average over Low and Intermediate activation levels) decreases with combinations of shorter recovery times and higher conditioning amplitudes (Fig. 9) characteristic of abnormal postural sway. This models indicated the presence of a plateau in the reduction of muscle short-range stiffness at combinations of conditioning amplitudes greater than 3% and recovery times lower than 0.5 s, characteristic of normal postural sway. Averaged over all experimental conditions (“All trials”, Fig. 10), relative short-range stiffness of the XB coop and XB coop + FD model deviated less from experimental values (relative SRS error = 0.011) compared with those from the XB model (relative SRS error = 0.017) and Hill-type model (relative SRS error = 0.031). Despite fitting most conditions better than the XB model (Fig. 8), the Hill-type model had a larger mean SRS error because of its large error at the lowest muscle activation (left-most data point in Fig 8A). Due to having fewer free parameters than the XB coop + FD model, the XB coop model yielded a lower ΔAIC score (ΔAIC = −80 versus −76). Both ΔAIC scores were considerably lower than those from the XB coop model (ΔAIC = −42), indicating greater model quality. Averaged over trials with little to no history-dependent short-range stiffness changes (“History-independent trials”, Fig. 10), all biophysical models deviated a similar amount from experimental data (SRS error = 0.011-0.012). The Hill-type model deviated a bit more (SRS error = 0.023). Averaged over trials with large history-dependent short-range stiffness changes (“History-dependent trials”, Fig. 10), relative short-range stiffness of the XB coop + FD model deviated least from experimental values (SRS error = 0.006), followed by the XB coop model (SRS error = 0.008), the XB model (SRS error = 0.055) and the Hill-type model (SRS error = 0.076). Altogether, the XB coop model yields greatest accuracy (based on ΔAIC) across all conditions in the dataset, while the XB coop + FD model yields smallest short-range stiffness errors when only considering the history-dependent conditions.

**Figure 9.**
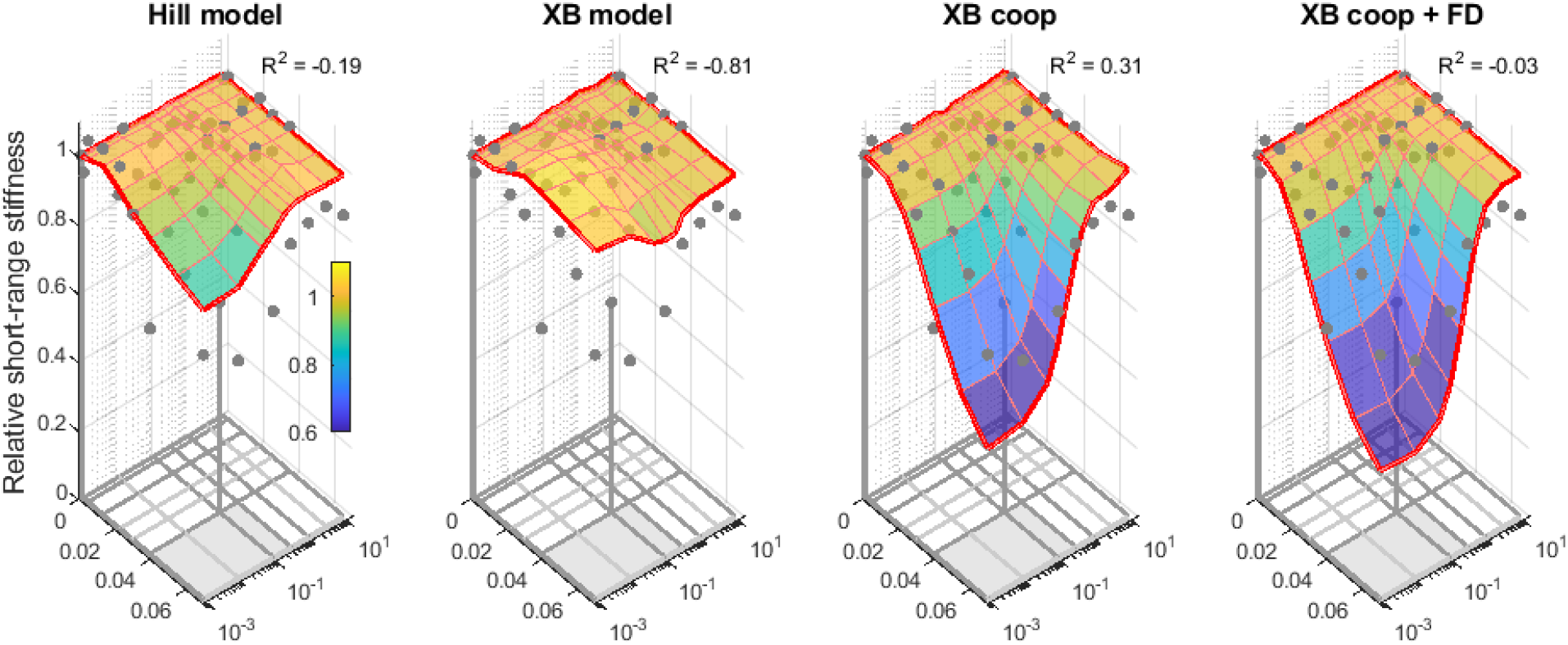
Empirical and model relative short-range stiffness at submaximal activation across stretch amplitudes and recovery times. Model predictions averaged over all fibers in the fitting set (red lines, n = 7) are shown for Hill-type model (“Hill model”), cross-bridge model (“XB model”), cross-bridge model with cooperative dynamics (“XB coop”), and cross-bridge model with both cooperative dynamics and a forcibly detached state (“XB coop + FD”). Relative short-range stiffness is indicated by both the vertical location and surface color. Experimental data are averaged across fibers for which specific conditions were collected (grey dots). Fitting trials are indicated with stems, connecting to the corresponding dots. The bottom surface indicates the experimental grid of stretch amplitudes and recovery times, distinguishing between conditions measured in all fibers (dark grey lines, n = 11) and conditions measured in a subset of fibers (light grey lines, n = 3), conditions with history-dependent stiffness reductions (grey surface) and conditions without such stiffness reductions (white surface).

**Figure 10.**
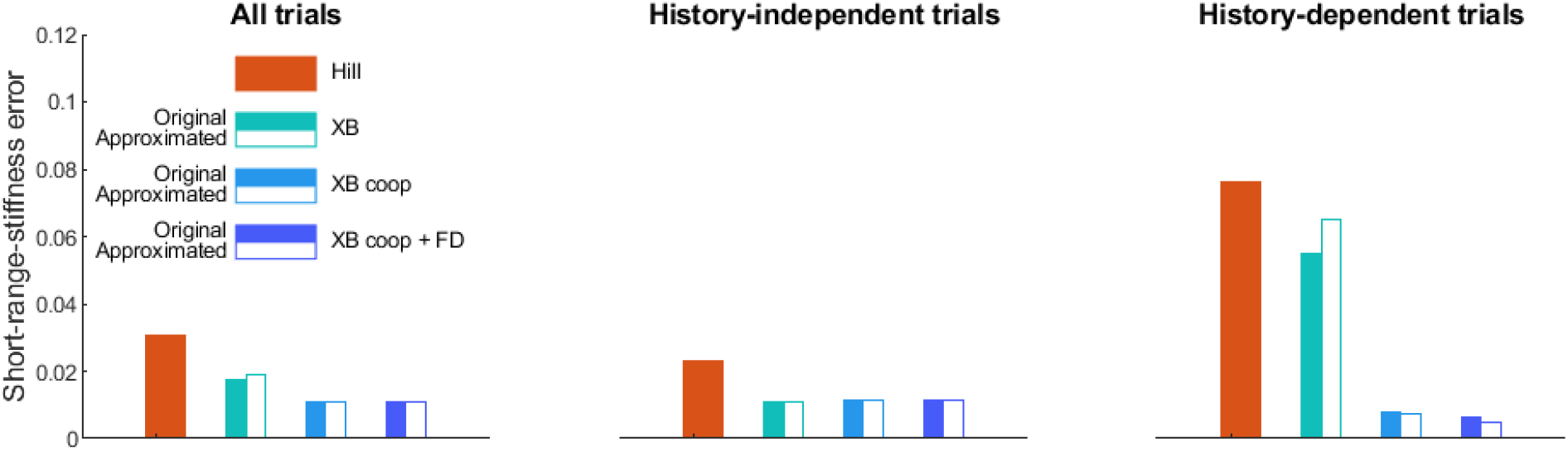
Short-range stiffness error for all trials, history-independent trials, and history-dependent trials.

**Approximating the cross-bridge distribution with a Gaussian has a small effect on predicted force trajectories during stretch-shortening but greatly reduced the computational demands.** The two solution methods for simulating muscle dynamics based on biophysical muscle models, i.e. discretizing the cross-bridge distribution and approximating the cross-bridge distribution by a Gaussian, produced similar force trajectories. Using a discretized instead of an approximated solution method had a small effect on force trajectories (Fig. 4-6 versus Fig. S1-3), RMSDs (Fig. 7 versus Fig. S4) and short-range stiffness predictions (Fig. 8-9 versus Fig. S5-6). As a result, short-range stiffness errors of discretized and approximated models were similar (closed versus open bars in Fig. 10). Compared to using a discretized solution method, using an approximated solution method typically yielded a ∼20 times shorter computation time (0.26 s versus 5.71 s, modern laptop with 11th Gen Intel(R) Core(TM) i7-1165G7 @ 2.80GHz) for simulating an entire trial (∼10 s). For a single state derivative function evaluation, processing time was ∼3 times shorter when using an approximated solution method (0.028 ± 0.014 ms, mean ± s.d.) with respect to all function calls of a trial) compared to using a discretized solution method (0.082 ± 0.012 ms). Most of the computational cost difference was thus due to a reduction in the number of function evaluations due to the larger time steps enabled by a reduction in the stiffness of the state derivative equations. Approximating the cross-bridge distribution by a Gaussian has little influence on the simulated force trajectory but improves computational efficiency.

## Discussion

Here we present a biophysical muscle model capturing skeletal muscle’s history-dependent reduction in short-range stiffness, referred to as muscle thixotropy. Beyond classic actin-myosin interactions, incorporating cooperative dynamics for myofilament activation greatly slowed force and stiffness recovery rates at submaximal activation – key to improving predictions of history-dependent short-range stiffness reductions. Adding a forcibly-detached myosin state further increased the history-dependence of short-range stiffness a little, but not necessarily in better agreement with data. Further, the resulting cross-bridge distributions that determine muscle force were well-described by Gaussian approximations, reducing computational time by a factor of 20. This faster computation time facilitates integration of biophysical muscle models in musculoskeletal simulation to test the role of muscle short-range stiffness in movement. Musculoskeletal simulations requiring history-dependent short-range stiffness may be achieved using Gaussian-approximated biophysical models with cooperative dynamics.

**Cooperative myofilament dynamics yield an activation-dependent reduction in cross-bridge cycling rates after shortening, which may underly the activation-dependent movement history-dependence of short-range stiffness.** Modeling cooperative myofilament dynamics was important to predict the slower rate of force redevelopment during the isometric recovery period between two consecutive stretches at submaximal compared to maximal activation in agreement with experimental data (Fig. 5-6, panel B versus A for condition with long recovery time). The Hill-type and XB model, which lack cooperative dynamics, predict similar force rates for submaximal and maximal activations (Fig. 5-6). As a result, they overestimated force recovery at submaximal activation, and underestimated the reduction in short-range stiffness. Cooperative myofilament activation may slow down cross-bridge (re-)attachment, particularly at submaximal activation (Fitzsimons et al., 2001; Campbell, 2014), because at low [Ca]^2+^. This may be because low myofilament activation (e.g. due to muscle shortening) suppresses the rate of myofilament activation and thereby the rate of cross-bridge cycling. Over time, cross-bridges slowly attach and thereby activate thin and thick myofilaments, which facilitates cross-bridges attachment, which facilitates myofilament activation, etc. leading to increasing force rates. As a consequence, it may take several seconds to reach a steady-state force (see Fig. 1 in Fitzsimons et al., 2001). In contrast, high [Ca]^2+^ may jumpstart this process by rapidly activating the thin filament, such that steady-state force is reached about twice as fast (see Fig. 1 in Fitzsimons et al., 2001). Thus, cooperative myofilament activation implies that the initial rate of cross-bridge (re-)attachment is disproportionately slower at submaximal activation (Campbell, 2014). This mechanism may explain why we found that biophysical models with cooperative myofilament activation yield better predictions of history-dependent changes in short-range stiffness, particularly at submaximal activation (Figs. 8-9). Yet, force increases during lengthening after shortening were still somewhat overestimated (Fig. 4-6). Future modeling studies could explore whether modelling local thin and thick filament activation (Kosta et al., 2022; Liu et al., 2024; Walcott, 2014; Walcott et al., 2012) further improves force predictions.

**A forcibly detached cross-bridge state helps to reconcile fast cross-bridge cycling during stretch with slow recovery after shortening but the small improvements in short-range stiffness predictions do not justify the additional complexity.** While history-dependent short-range stiffness reduction requires slow cross-bridge cycling rates, force transients during lengthening require fast cycling rates. Cooperative dynamics partially resolve this, as they explain why cross-bridge cycling rates are temporally reduced after shortening and can increase again during lengthening. However, there remains a trade-off between slow and fast dynamics. This may explain why cooperative XB models still slightly underestimate short-range stiffness reductions (light blue lines, Fig. 8). Adding a forcibly detached state partially resolves this. While a forcible detached state is only entered at large strains that do not occur within the short-range stiffness time interval, parameters are estimated based on entire force trajectories and are thus a trade-off between fitting the force plateau during stretch requiring fast dynamics and the recovery requiring slow dynamics. Forcible detachment dynamics are fast and explain forces during the later stages of stretch, while regular detachment dynamics are slow (due to cooperative dynamics) and explain short-range stiffness reductions. Despite this advantage, the additional complexity and number of parameters did not justify the inclusion of a forcibly detached state to describe history-dependent short-range stiffness (based on AIC). Evidence for the existence of a forcibly detached state is much weaker than evidence for cooperative myofilament dynamics. Instead of forcible detachment, a weakly-bound cross-bridge state may be used. A weakly-bound cross-bridge state with fast dynamics has been suggested as an alternative mechanism to explain the force plateau during stretch. There is more evidence for such weakly bound state than for forcible detachment (Huxley and Simmons, 1971; Campbell and Moss, 2000; Campbell and Moss, 2002; Lakie and Campbell, 2019). It should be explored whether such models with a weakly bound state can improve predictions of the force plateau during stretch and thereby the overall agreement with experimental force trajectories.

**A Gaussian approximation provides a good trade-off between accuracy and computational demand.** Force traces simulated based on a discretization of the cross-bridge distribution, which does not require any assumptions about its shape, were similar to force traces simulated under the assumption of a Gaussian distribution, yet computational cost was ∼20 times smaller for the approximated model. In particular, errors in the prediction of short-range stiffness were similar when using discretization or a Gaussian approximation to simulate cross-bridge dynamics (Fig. 10). Zahalak (1981) was the first to approximate the cross-bridge distribution with a Gaussian, arguing that although the actual cross-bridge distribution may not be Gaussian, a Gaussian function may adequately reflect the distribution’s first three moments. Conveniently, these moments have physical representations, namely the ensemble cross-bridge stiffness, force, and elastic strain energy. While Zahalak (1981) used Huxley’s (1957) piece-wise linear cross-bridge attachment function (proposed for mathematical convenience rather than mechanistic accuracy), more recent findings indicate that a Gaussian attachment better captures the underlying mechanisms. This is because detached myosin heads are thought to have random Brownian motion that is described by a Gaussian. More recent cross-bridge models (e.g., Campbell et al., 2018) therefore use a Gaussian attachment function, yielding a Gaussian-like cross-bridge distribution during isometric muscle contraction. In addition, unlike Zahalak’s and Huxley’s piece-wise linear detachment functions, cross-bridge detachment functions are thought to be exponentials, which help preserve a Gaussian-shaped distribution during length changes. Thus, a Gaussian may be an even better approximation than originally conceived by Zahalak. In line with this idea, we found that biophysical models with discretized (Figs. 4-8) and approximated solution methods (Figs. S1-S5) predicted similar forces during stretch-shortening. This finding is also in line with a recent study that found good agreement between approximated and discretized solution methods during force development (van der Zee et al., 2024). The smooth shape of Gaussian and exponential attachment and detachment functions also enabled a lower bin resolution and thereby far fewer bins in the discretized model compared with models that use piece-wise linear rate functions (van Soest et al., 2019), decreasing the computational cost per iteration. The bin resolution used here (0.71 nm) is similar to that of other models using a Gaussian attachment function (e.g., 0.5 nm, Campbell et al., 2018). Smoother attachment and detachment functions also resulted in differential equations that were less stiff, thereby decreasing the number of required time steps. Nevertheless, the reduced number of integration steps needed when using a Gaussian approximation suggest that the dynamics of the discretized model were stiffer than those of the approximated model, contributing about two-thirds to the overall increased computational demand.

**We focused on history-dependent short-range stiffness here but cooperative myofilament dynamics may also contribute to other aspects of muscle mechanics.** While our simulations suggest that cooperative dynamics may underly muscle thixotropy, these dynamics can also explain other aspects of muscle mechanics. These include the sigmoidal relation between steady-state isometric force and calcium concentration (Campbell et al., 2018) and ‘muscle’s low-pass filter properties in response to cyclic excitation (van der Zee et al., 2024). It is possible that cooperative myofilament dynamics also contributes to aspects of history dependence of muscle force production beyond short-range stiffness, such as residual force depression and residual force enhancement (Abbott and Aubert, 1952). A previous biophysical muscle model has suggested a role for cooperative myofilament dynamics in residual force depression, i.e. a reduction in muscle force after shortening (Liu et al., 2024). Residual force enhancement is an increase in muscle force after stretch, thought to be mediated by titin (Schappacher-Tilp et al., 2015; Liu et al., 2024; Nishikawa et al., 2012) although others have suggested that it may also arise from half-sarcomere inhomogeneity mediated by cross-bridge cycling (Campbell, 2009; Campbell et al., 2011). One theory is that cross-bridge cycling strains titin and thereby increases force produced by titin (Nishikawa et al., 2012). If this is the case, cooperative myofilament dynamics will also influence the rate at which titin is strained and thus residual force enhancement. Further, it has been suggested that titin could contribute to myosin activation as it is connected to and can thus transfer force to myosin (Squarci et al., 2023). It is difficult to isolate the effects of individual myofilament dynamics on force production. To date, biophysical muscle models have generally either included titin dynamics (Schappacher-Tilp et al., 2015) or cooperative myofilament dynamics (Campbell et al., 2018; Liu et al., 2024) but not both. Residual force enhancement has been explained by models lacking titin dynamics (Campbell, 2009; Campbell et al., 2011) but as far as we know, no existing biophysical model has simultaneously explained all aspects of muscle history dependence, including muscle thixotropy, residual force depression and residual force enhancement. Considering that these various aspects of muscle’s history dependence may arise from interactions between myosin, actine, and titin dynamics, future biophysical muscle models should integrate these dynamics to generate a comprehensive understanding of muscle history dependence.

**Modeling cooperative myofilament dynamics may be crucial to simulate whole-body movements.** Muscle thixotropy has been observed across a wide range of scales, ranging from single fiber to human movement (Lakie and Campbell, 2019) as well as in intrafusal muscle fibers encapsulating muscle spindles (Blum et al., 2020; Proske et al., 1993; Simha and Ting, 2023). Our findings might therefore have broad implications for understanding neuromechanics of whole-body movements. Simulations are a useful tool to disentangle the relation between muscle mechanisms and whole-body behavior, yet most simulations of whole-body movement are based on Hill-type models and the few simulations that are based on cross-bridge dynamics lack cooperative myofilament dynamics (Wakeling et al., 2023). Some simulations have used phenomenological models of short-range stiffness that require the user to explicitly define the movement history and do not capture the complex relationships between pre-stretch characteristics, activation and short-range stiffness (De Groote et al., 2017; Willaert et al., 2024). These simulations have shown that modeling the large short-range stiffness in muscles that have been isometric is crucial to explain the response to perturbations of standing (De Groote et al., 2017; Jakubowski et al., 2025; Loram et al., 2009; Van Wouwe et al., 2022), walking (Araz et al., 2023) and reaching (Hu et al., 2011). Furthermore, capturing the history dependence of short-range stiffness is key to explaining joint hyper-resistance in neurological populations (Groote et al., 2018; Willaert et al., 2020; Willaert et al., 2024). Yet, understanding the role of short-range stiffness during continuous or cyclic movement such as locomotion requires a model of muscle dynamics that captures how short-range stiffness varies with movement history and activation. The cross-bridge model with cooperative myofilament dynamics proposed here could fill this gap, especially since describing the cross-bridge distribution by a Gaussian yields high accuracy at low computational demand. The logical next step is therefore to implement these models in whole-body musculoskeletal simulations to simulate short-range stiffness (reductions) during movement.

## Conclusion

This study aimed to identify a muscle model based on its ability to capture skeletal muscle’s history-dependent reduction in short-range stiffness, referred to as muscle thixotropy. We found that incorporating cooperative dynamics for myofilament activation greatly slowed force and stiffness recovery rates at submaximal activation, which was key to improving predictions of history-dependent short-range stiffness reductions. Adding a forcibly-detached state yielded somewhat greater history-dependent reductions in short-range stiffness, but not necessarily in better agreement with data. Using a Gaussian approximation of the cross-bridge distribution seems to be a good trade-off between model accuracy and computational demand, facilitating integration in musculoskeletal simulation. Such simulations could elucidate the role of history-dependent short-range stiffness in whole body movement.

## Code availability

All code pertaining to the muscle models is available on GitHub (https://github.com/timvanderzee/biophysical-muscle-model).

## Appendix

Analytical expressions for the spatial integrals

Because the attachment functions *f*(*x*) and *ϕ*(*x*) are gaussian, *C*_*λ*_ and *β*_*λ*_ can be expressed in terms of model parameters *f*_1_, *ϕ*_1_ and *w*:

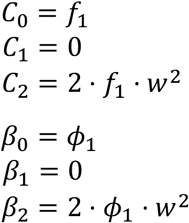

Because an exponential detachment function *g*(*x*) is used, *θ*_*λ*_ can be written in terms of the moments of the cross-bridge distribution *Q*_0_ - *Q*_2_, and the model parameters *g*_1_, *g*_2_, *E*_1_ and *E*_2_:

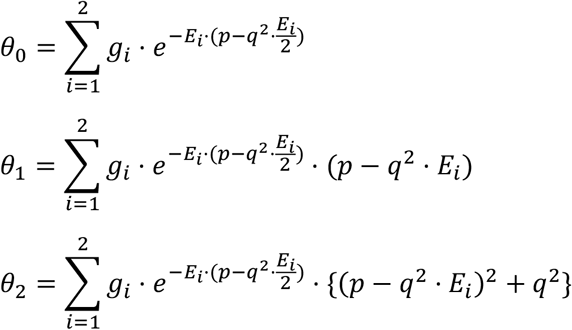

Because the detachment function Γ(*x*) is a step function, Ω_*λ*_ can be written in terms of the moments of the cross-bridge distribution *Q*_0_ - *Q*_2_, and the model parameters *r* and *x*_*c*_:

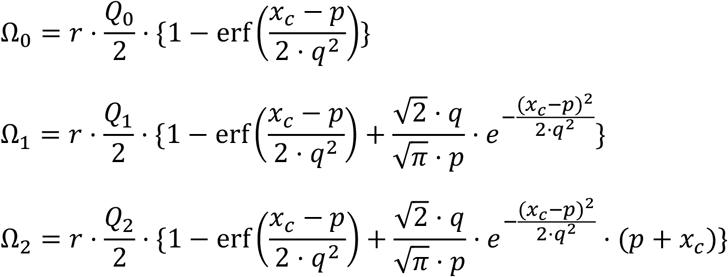

Here, erf is the error function.

## Supplementary figures

**Figure S1.**
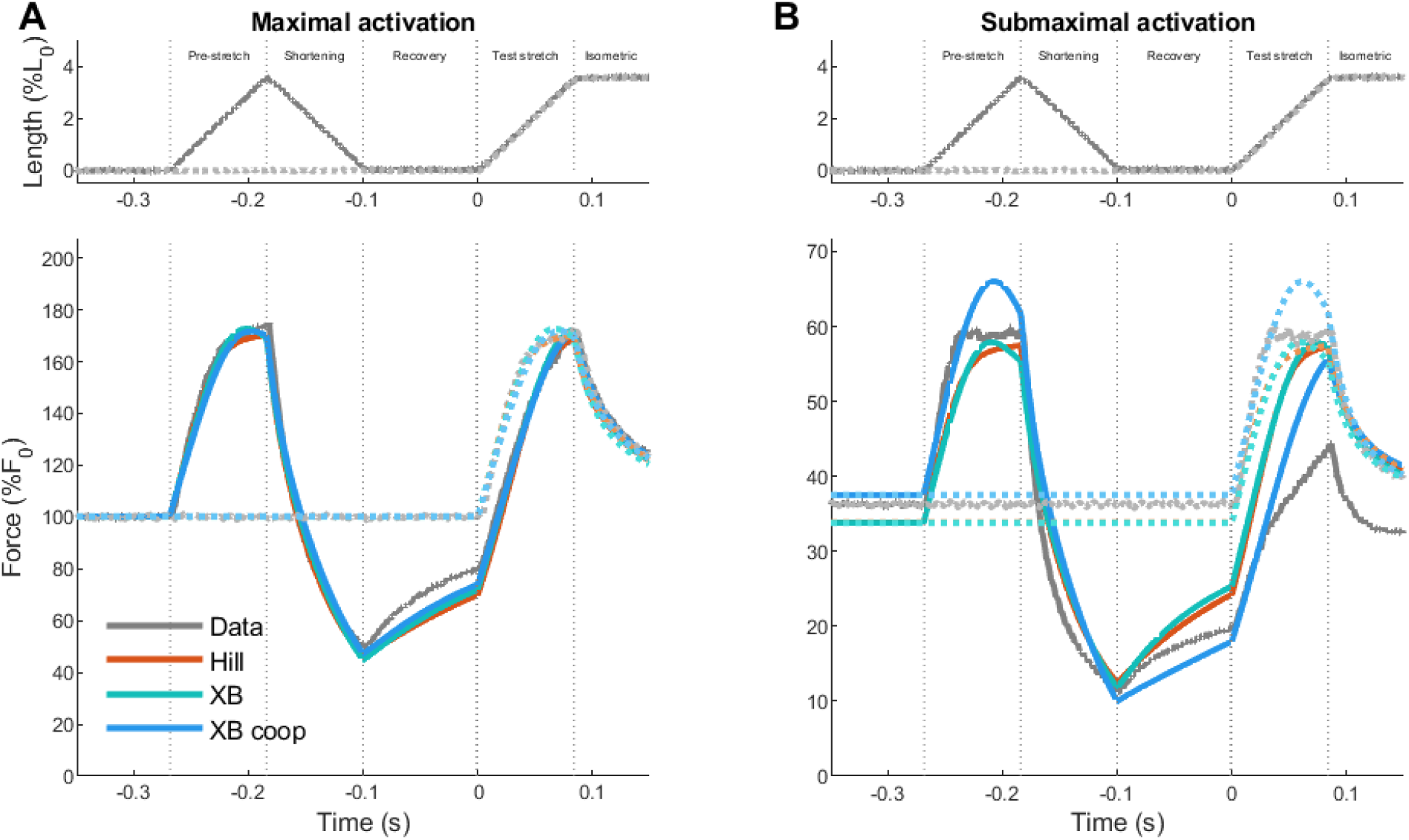
Approximated modelled and measured forces for fitting trials: effect of cooperative dynamics. Top row shows measured fiber length changes for a condition with a pre-stretch (darker solid lines) and without a pre-stretch (lighter dotted lines). Bottom row shows the corresponding empirical forces (“Data”, grey), alongside forces of Hill-type model (“Hill”, red), cross-bridge model (“XB”, green), cross-bridge model with cooperative dynamics (“XB coop”, blue). **A**: maximal activation. **B**: submaximal activation.

**Figure S2.**
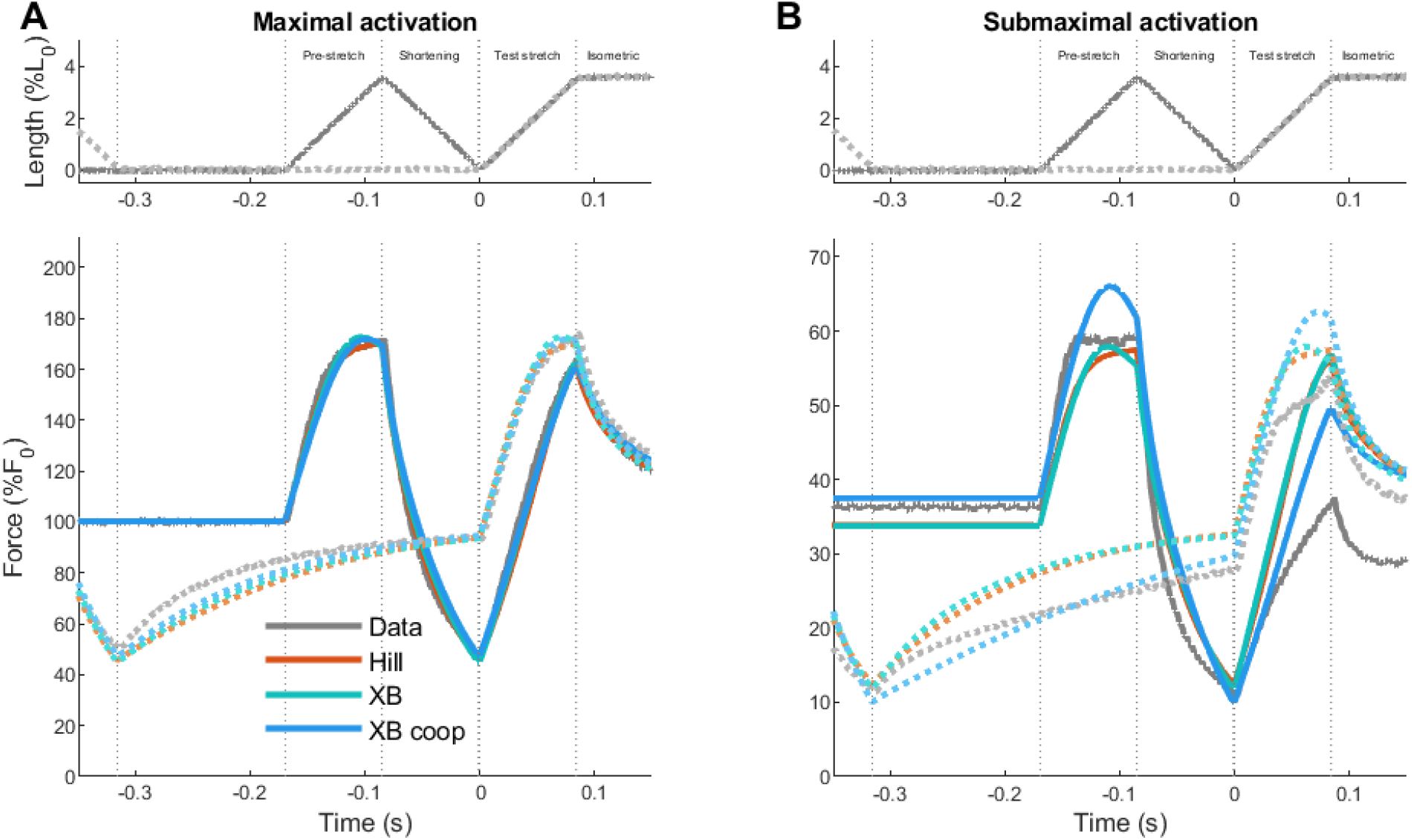
Approximated modelled and measured forces for two testing trials: effect of cooperative dynamics. Top row shows measured fiber length changes for a condition with short recovery time (darker solid lines) and long recovery time (lighter dotted lines). Bottom row shows the corresponding empirical forces (“Data”, grey), alongside forces of Hill-type model (“Hill”, red), cross-bridge model (“XB”, green), cross-bridge model with cooperative dynamics (“XB coop”, blue). **A**: maximal activation. **B**: submaximal activation.

**Figure S3.**
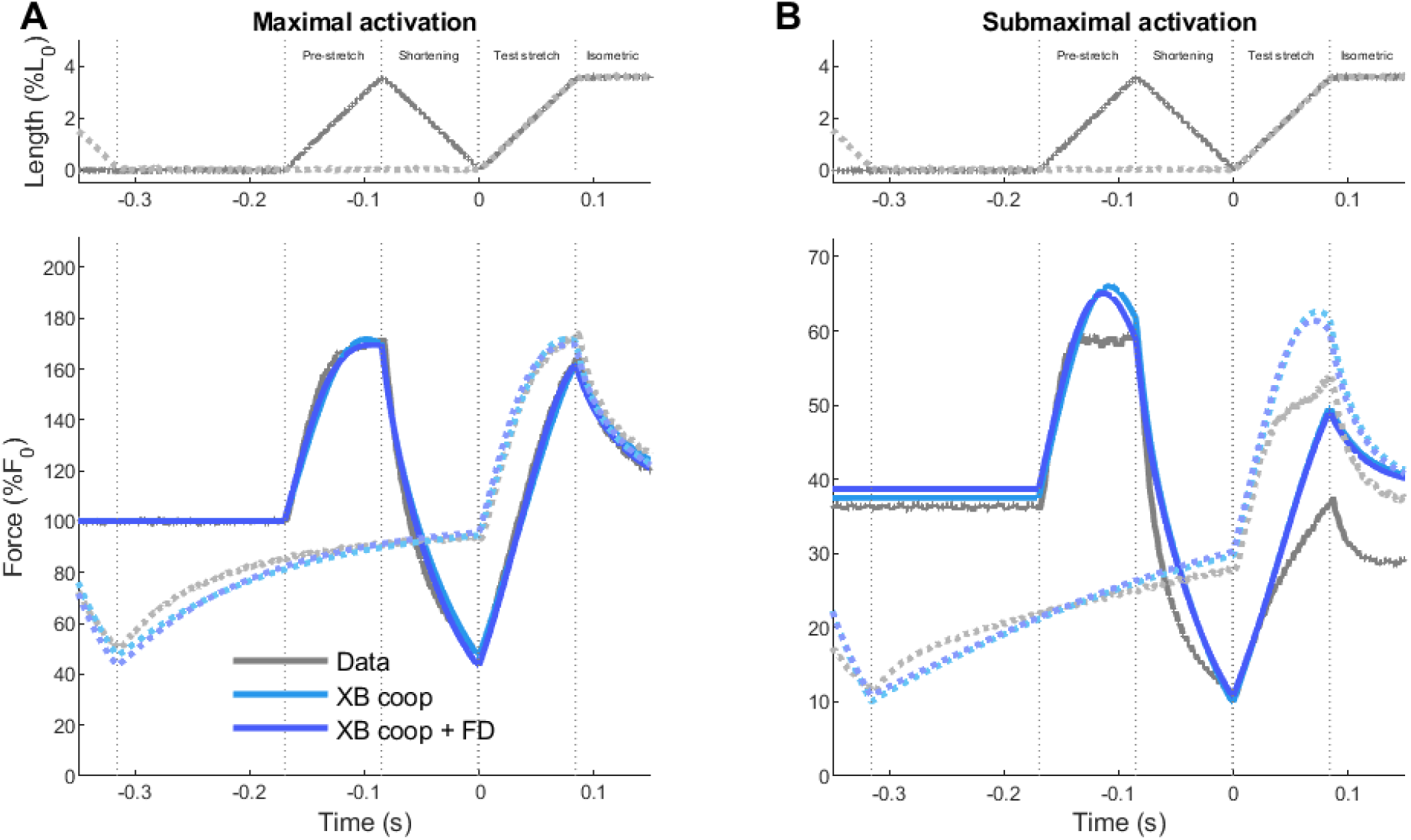
Approximated modelled and measured forces for two testing trials: effect of forcibly detached state. Top row shows measured fiber length changes for a condition with short recovery time (darker solid lines) and long recovery time (lighter dotted lines). Bottom row shows the corresponding empirical forces (“Data”, grey), alongside forces of Hill-type model (“Hill”, red), cross-bridge model (“XB”, green), cross-bridge model with cooperative dynamics (“XB coop”, light blue), and cross-bridge model with cooperative dynamics and a forcibly detached state (“XB coop + FD”, dark blue). **A**: maximal activation. **B**: submaximal activation

**Figure S4.**
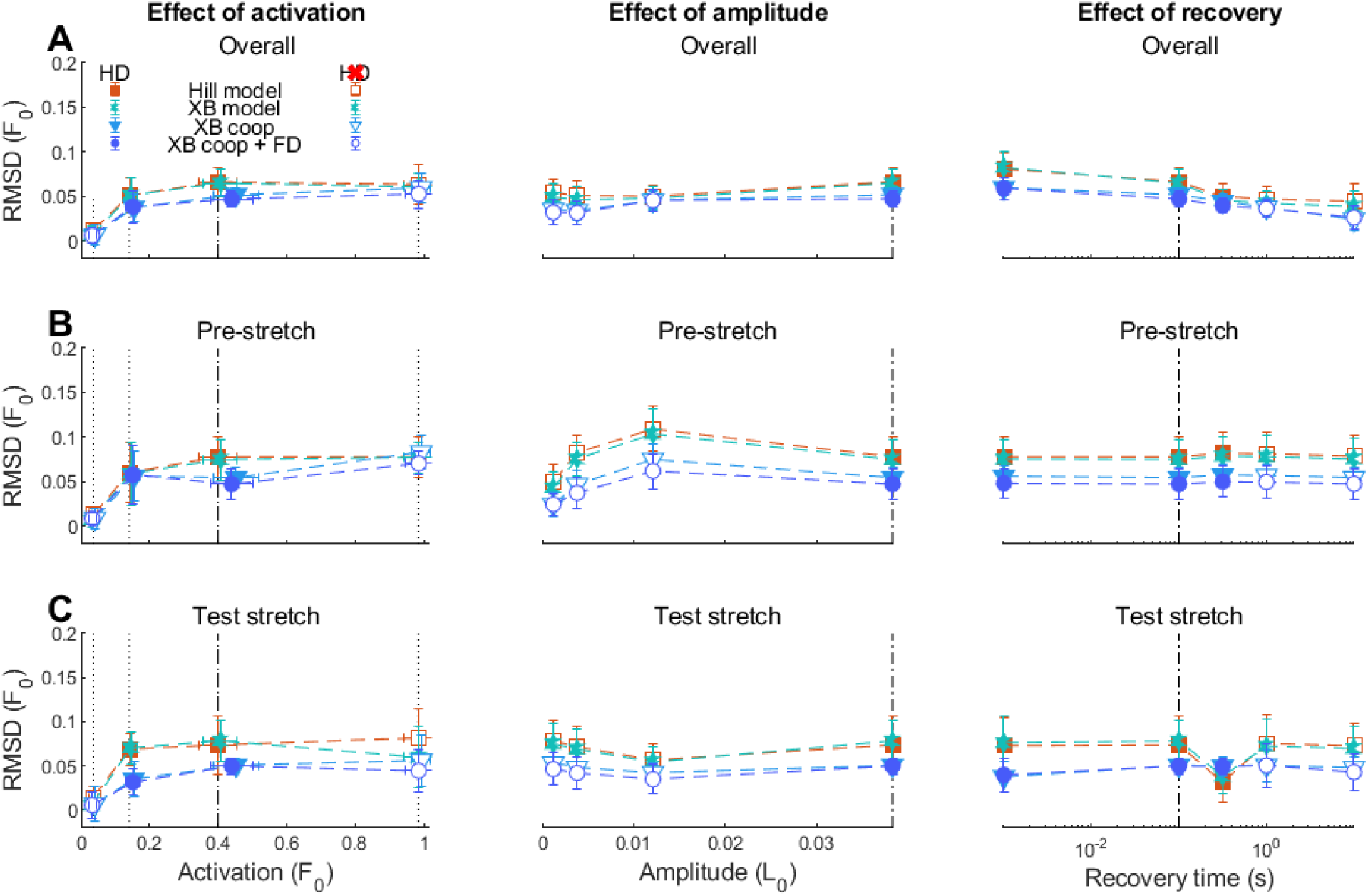
Root-mean-squared deviations between approximated model forces and experimental forces. RMSDs are shown for Hill-type model (“Hill”, red), cross-bridge model (“XB”, green), cross-bridge model with cooperative dynamics (“XB coop”, blue), and cross-bridge model with both cooperative dynamics and a forcibly detached state (“XB coop + FD”, dark blue). Filled symbols indicate trials with considerable short-range stiffness reductions (i.e. history dependence), open symbols indicate trials with little to no short-range stiffness reductions (i.e. no history dependence). Left column: effect of activation at constant amplitude and recovery time. Right column: effect of amplitude at constant activation and recovery time. Right column: effect of recovery time at constant activation and amplitude. Dashed vertical lines indicate the condition common to all three sets of conditions, dotted vertical lines indicate fitting trials (dashed-dotted indicate both). **A**: Entire stretch-shortening protocol. **B**: Pre-stretch. **C**: Test stretch.

**Figure S5.**
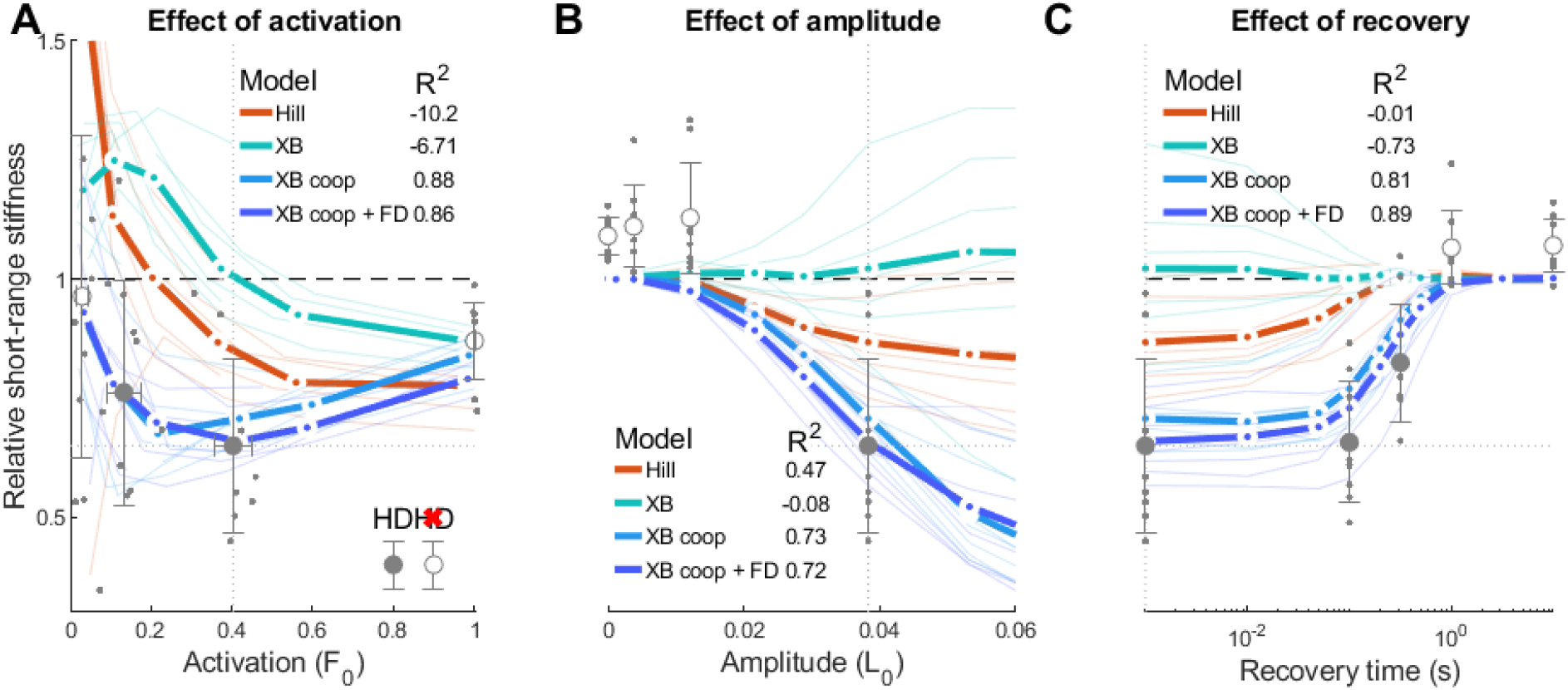
Empirical and approximated model relative short-range stiffness across muscle activations, stretch amplitudes and recovery times. Dotted lines indicate the condition common to all three sets of conditions. Model predictions are shown for Hill-type model (“Hill”, red), cross-bridge model (“XB”, green), cross-bridge model with cooperative dynamics (“XB coop”, blue), and cross-bridge model with both cooperative dynamics and a forcibly detached state (“XB coop + FD”, dark blue). Model predictions are shown for fibers in the fitting set (n = 7), both averaged across these fibers (thick lines) and for individual fibers (thin lines). Data is shown both averaged across all fibers (n = 11, grey circles), and for individual fibers (grey dots). Filled circles indicate trials with considerable short-range stiffness reductions (i.e. history dependence, HD), open symbols indicate trials with little to no short-range stiffness reductions (i.e. no history dependence, no HD). **A**: Effect of activation at constant amplitude and recovery time. **B**: Effect of amplitude at constant activation and recovery time. **C**: Effect of recovery time at constant activation and amplitude.

**Figure S6.**
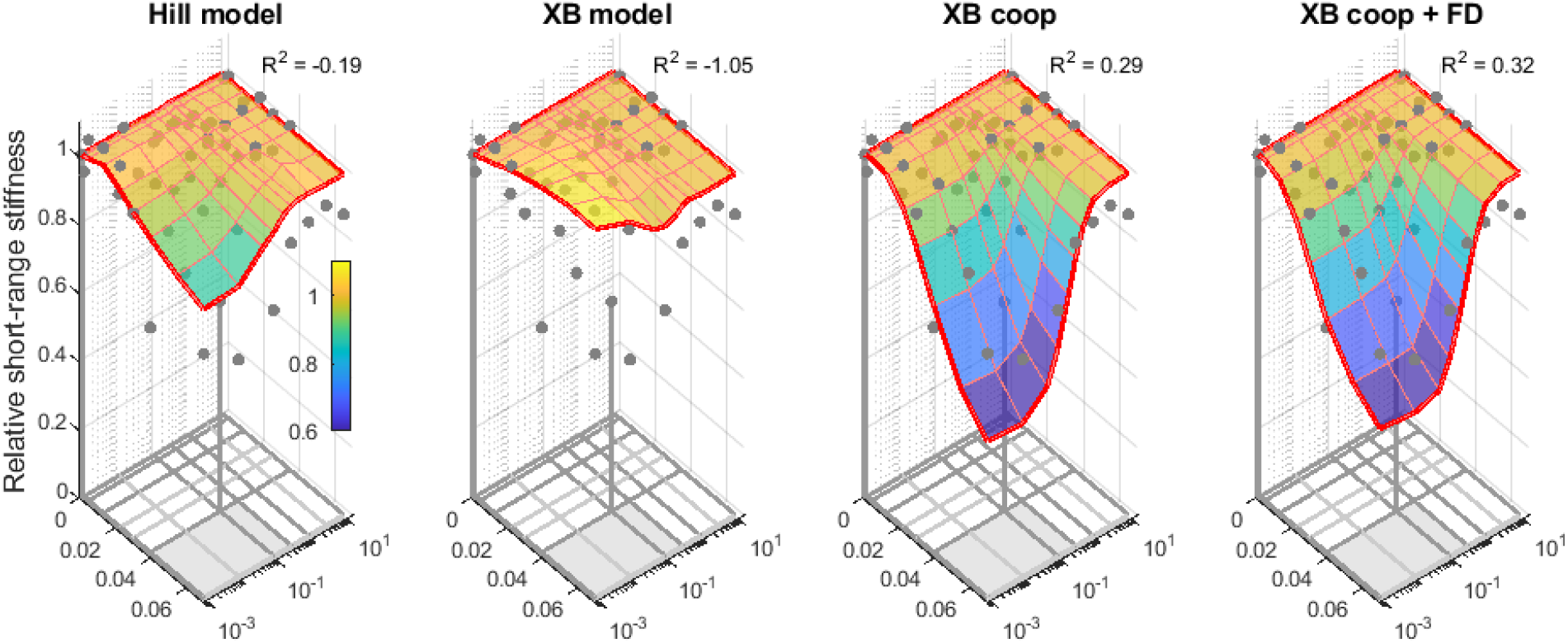
Empirical and approximated model relative short-range stiffness at submaximal activation across stretch amplitudes and recovery times. Model predictions averaged over all fibers in the fitting set (red lines, n = 7) are shown for Hill-type model (“Hill model”), cross-bridge model (“XB model”), cross-bridge model with cooperative dynamics (“XB coop”), and cross-bridge model with both cooperative dynamics and a forcibly detached state (“XB coop + FD”). Relative short-range stiffness is indicated by both the vertical location and surface color. Experimental data are averaged across fibers for which specific conditions were collected (grey dots). Fitting trials are indicated with stems, connecting to the corresponding dots. The bottom surface indicates the experimental grid of stretch amplitudes and recovery times, distinguishing between conditions measured in all fibers (dark grey lines, n = 11) and conditions measured in a subset of fibers (light grey lines, n = 3), conditions with history-dependent stiffness reductions (grey surface) and conditions without such stiffness reductions (white surface).

